# Redefining paradigms of cellular HPV trafficking *in situ*

**DOI:** 10.1101/2025.09.06.674660

**Authors:** Renaldo Sutanto, Jaimin K. Rana, Chao-Yin Cheng, Paul Lephart, Diane M. Harper, Billy Tsai, Shyamal Mosalaganti

## Abstract

Human papillomavirus (HPV) is the most common sexually transmitted agent and causes ∼4.5% of all human cancers. A complete understanding of the cellular mechanisms underlying trafficking, disassembly, and transport is still lacking. Using *in situ* cryo-electron tomography, we directly visualize high-risk HPV16 during infection and reveal that virions remain intact and infection-competent upon reaching lysosomes, contrary to the prevailing model of lysosomal inactivation. We further challenge the established view that the L1 capsid progressively disassembles during retrograde transport by showing that HPV can remain intact upon reaching the Golgi *en route* to the nucleus. Finally, we provide snapshots of HPV-containing transport vesicles budding from the Golgi lumen in a COPI coat-dependent fashion and show that HPV is connected to the overlying membrane by proteinaceous bridges. Together, these results challenge current models of HPV entry by revealing the capsid’s resilience to disassembly during retrograde trafficking, identifying a coat-dependent budding mechanism, and demonstrating that lysosomes harbor a persistent pool of intact HPV particles that retain infectivity.

## Introduction

Human papillomaviruses (HPV) are a group of >200 related viruses, over 40 of which infect the anogenital and/or oropharyngeal mucosa, and are sexually transmitted ^1–3^. Infection is highly prevalent; however, 90% of infections clear spontaneously within ∼2 years without symptoms ^4,5^, but persistent infection with oncogenic (high-risk) types (e.g., HPV16/18) can lead to precancerous lesions and cancer ^3,6^. Globally, HPV accounts for ∼4.5% of all cancers ^7,8^. It is the leading cause of cervical cancer and contributes significantly to anogenital and oropharyngeal cancers ^9–11^. HPV also causes a wide range of warts, including common warts, plantar warts, and anogenital warts ^12,13^. Although effective prophylactic vaccines against multiple high-risk HPV types are available, there are currently no antiviral drugs, and the global vaccine coverage remains less than 15% ^14,15^.

Structurally, HPV is a non-enveloped DNA tumor virus composed of 72 pentamers of the major capsid protein L1, which forms the outer shell of the capsid ^16–22^, and up to 72 copies of the minor capsid L2 protein buried within the capsid shell ^20–22^. When fully assembled, the capsid is approximately 55 nm in diameter ^16–22^ and encapsulates an 8 kb viral genome ^23^. To initiate cell entry, the L1 capsid binds to heparan sulfate proteoglycans (HSPGs) on the host plasma membrane, enabling furin-mediated cleavage of L2 at the cell surface ^21,24–28^. Following cleavage, the virus is transferred to cell-surface receptor(s) and internalized via actin-mediated endocytosis ^29,30^. The acidic endosomal environment is proposed to trigger partial disassembly of the HPV L1 capsid, exposing the C-terminus of the L2 protein ^31–44^. The cell-penetrating peptide (CPP) motif at the extreme C-terminus of L2 ^45–47^, in concert with the host protein γ-secretase, assists in the insertion of L2 into and across the endosomal membrane ^43,48,49^. This exposes a portion of L2 to the cytosol, enabling L2 to recruit cellular trafficking factors—including retromer, retriever, and the dynein-BICD2 motor complex—that target HPV to the trans-Golgi network (TGN) ^50–58^.

Upon reaching the TGN ^59^, the cytosol-exposed segment of L2 binds directly to the coat protein I (COPI) coatomer complex ^60^. This coat protein complex is postulated to induce the formation of membrane-bound vesicles that ferry HPV from the TGN and across the Golgi stacks. When HPV reaches the *cis*-Golgi compartment, importin receptors, including IPO7, help deliver HPV to the nuclear membrane ^61^. Nuclear envelope breakdown at the onset of mitosis enables the virus to enter the nucleus and access the genome to cause infection ^62–64^. Therefore, retrograde trafficking along the endosome-TGN/Golgi-nucleus axis is thought to represent a productive infection route, although whether there exist other pathways leading to infection remain unknown. By contrast, because endosome-localized HPV can also be delivered to the lysosome where it is thought to be degraded, endosome-to-lysosome transport of HPV has been considered a non-productive route ^30,32–34,38,65,66^.

Despite its clinical importance, HPV entry, intracellular trafficking, and uncoating remain incompletely understood. A central question is the extent of HPV disassembly during retrograde trafficking, as biochemical studies suggest a progressive, stepwise disassembly ^51,67–70^; however, visual evidence to support this model is lacking. The identity of transport vesicles mediating retrograde HPV movement is also unclear. Disruption of host factors involved in membrane tubulation and vesicle formation (including retromer ^47,51,53,56,57^, sorting nexins ^50,54,55^, and COPI ^60^) blocks infection, highlighting their importance in virus transport. However, due to the asynchronous nature of HPV entry and the transient presence of the virus in these compartments, direct high-resolution visualization of the virus in membrane-bound vesicles has not been achieved ^71^.

Here, using the high-risk HPV16 model system, we redefine the prevailing model of HPV retrograde trafficking by providing unprecedented, high-resolution *in situ* snapshots of key entry events. Integrating cryogenic correlative light and electron microscopy (cryo-CLEM) with *in situ* cryo*-*electron tomography (cryo-ET), pharmacological perturbations, confocal microscopy, and biochemical assays, we demonstrate that: (i) contrary to the canonical view that lysosomes degrade HPV, virions within the lysosomal compartment remain structurally intact and infectious; (ii) intact HPV is retrograde trafficked to the Golgi; (iii) intact HPV is a cargo of COPI-coated vesicles by providing high-resolution cellular snapshots; and (iv) the vesicular carriers that ferry HPV, and persist through mitosis, exhibit defined architectural features, with proteinaceous densities bridging the viral capsid and the membranes of Golgi compartments and transport vesicles. Together, these findings reposition endolysosomes as a reservoir and staging site for infectious HPV, establishing a revised framework for HPV trafficking.

## Results

### Characterizing fluorescently labeled HPV pseudovirus

To precisely localize HPV in the cell during infection, we first optimized a method to fluorescently label the well-established model system for studying papillomavirus infection mechanisms, the HPV16 pseudovirus (PsV) ^29,72^. These PsVs are composed of high-risk HPV16 L1 and L2 capsid proteins (where L2 is fused with a C-terminal 3x-FLAG tag) that encapsulate a GFP-S reporter plasmid in place of the native HPV16 viral genome. In this system, GFP-S (hereafter referred to as GFP) expression indicates successful delivery of PsV DNA to the nucleus, confirming productive entry into the cell. Importantly, HPV16 PsV infection closely mimics the entry pathway of authentic HPV isolated from stratified keratinocyte raft cultures; thus, this system has been widely adopted and used as a standard tool for investigating HPV entry mechanisms ^73–77^.

To generate a fluorescently labeled virus, purified HPV16 PsV (hereafter, HPV) was covalently labeled with Alexa Fluor 647 (**Figure S1A**: AF647-HPV). Successful fluorescent labeling was confirmed by SDS-PAGE, which revealed distinct fluorescent bands corresponding to the L1 and L2-FLAG proteins (**Figure S1B**). We verified that the AF647-HPV retained its structural integrity by negative staining electron microscopy (nsEM). AF647-HPV had an average diameter of 56.71 ± 2.13 nm (**Figure S1C**), consistent with the size of unlabeled HPV PsV ^16–22^. We next verified whether labeling the virus affected its infection competency. Using the widely studied cellular model for HPV trafficking, the immortalized cervical cancer cell line HeLa ^78^, we observed no significant difference in GFP expression between cells infected with labeled or unlabeled pseudovirus (**Figure S1D**; relative infection is quantified in the bottom graph). Infection by AF647-HPV was also sensitive to pharmacological treatment that disrupts critical HPV infection steps: XXI, a γ-secretase inhibitor that blocks L2 insertion across the endosomal membrane required for trafficking ^48,79–81^; NAV-2729, an Arf1 inhibitor that interferes with COPI recruitment to the Golgi necessary for retrograde transport across the Golgi ^60,82,83^; and RO-3306, a Cdk1 inhibitor, which prevents nuclear envelope breakdown essential for HPV nuclear entry (**Figure S1E**; normalized infection is quantified in the bottom graph) ^84,85^. Additionally, at 22 hours post-infection (h.p.i), when HPV begins to reach the Golgi, AF647-HPV colocalizes strongly with the Golgi marker HA-COPγ1-mCherry (**Figure S1F**), with quantitative analysis demonstrating a high Pearson Correlation Coefficient (PCC) between AF647-HPV, L1, and HA-COPγ1-mCherry (**Figure S1G-I**). Finally, to determine whether labeling altered trafficking or its timing, we compared colocalization of labeled and unlabeled virions with two compartments at well-established time points: the early endosome marker EEA1 at 4 h.p.i. (**Figure S1J**) and the trans-Golgi network marker TGN46 at 16 h.p.i (**Figure S1K**) ^30,50,59^. We observed no significant differences in the PCC between the two preparations, indicating that AF647 labeling does not significantly alter the timing of HPV trafficking.

Together, these findings demonstrate that AF647-HPV uses the same entry pathway as the unlabeled virus to promote infection, supporting its use as a reliable tool for detailed imaging and mechanistic studies of virus trafficking.

### Lysosomes harbor a persistent pool of infection-competent HPV particles

The canonical model of HPV trafficking posits that, after reaching the endosome from the cell surface, HPV is targeted either along the TGN/Golgi axis for productive infection or to the lysosome for degradation, rendering it ineffective for infection. To examine this paradigm, we first determined whether HPV is present in lysosomes using confocal microscopy. We found that HPV L1 colocalizes strongly with the lysosomal marker LAMP1 as early as 8 h.p.i (**Figure S2A,** 1^st^ row; quantified in **Figure S2B**), consistent with previous reports ^30,32–34,38,65,66^. The extent of L1-LAMP colocalization increases at 16 h.p.i (**Figure S2A**, 2^nd^ row; quantified in **Figure S2B**) and remains at this high level even at 72 h.p.i (**Figure S2A**, 3^rd^-5^th^ rows; quantified in **Figure S2B**).

Given our initial finding of HPV L1 retention within lysosomes, we questioned whether HPV remained intact within the acidic environment. To enable high-resolution *in situ* visualization, we employed a cryogenic-correlative light and electron microscopy workflow (cryo-CLEM, **Figure S3**), where HeLa cells were infected with AF647-HPV for 16 hours, treated with LysoTracker for 3-5 minutes to localize lysosomes, and plunge-frozen in liquid ethane. Cryo-confocal microscopy of these cells revealed near-perfect fluorescence colocalization between HPV and lysosomes, enabling precise cryo-focused ion beam (cryo-FIB) milling and cryo-electron tomography (cryo-ET) tilt series data collection (**Figures 1A and 1B)**. We observed that lysosomes contain varying numbers of intact, visually discernible HPV capsids (**Figure S2I and S2J**). In rare cases, lysosomes are densely packed with several HPV particles (**Figures 1C and 1D; Video S1**). Strikingly, these lysosomal HPV were indistinguishable from purified HPV PsV, retaining the characteristic appearance of the L1 pentamer (**Figure 1C i-iii**, yellow arrowheads), and contained a central density, likely plasmid DNA (**Figure 1C i-iii**, green arrowheads). Moreover, the diameter of lysosomal HPV particles was similar to that of purified HPV PsV (57.05 ± 1.96 nm) (**Figure 1E**).

**Figure 1.**
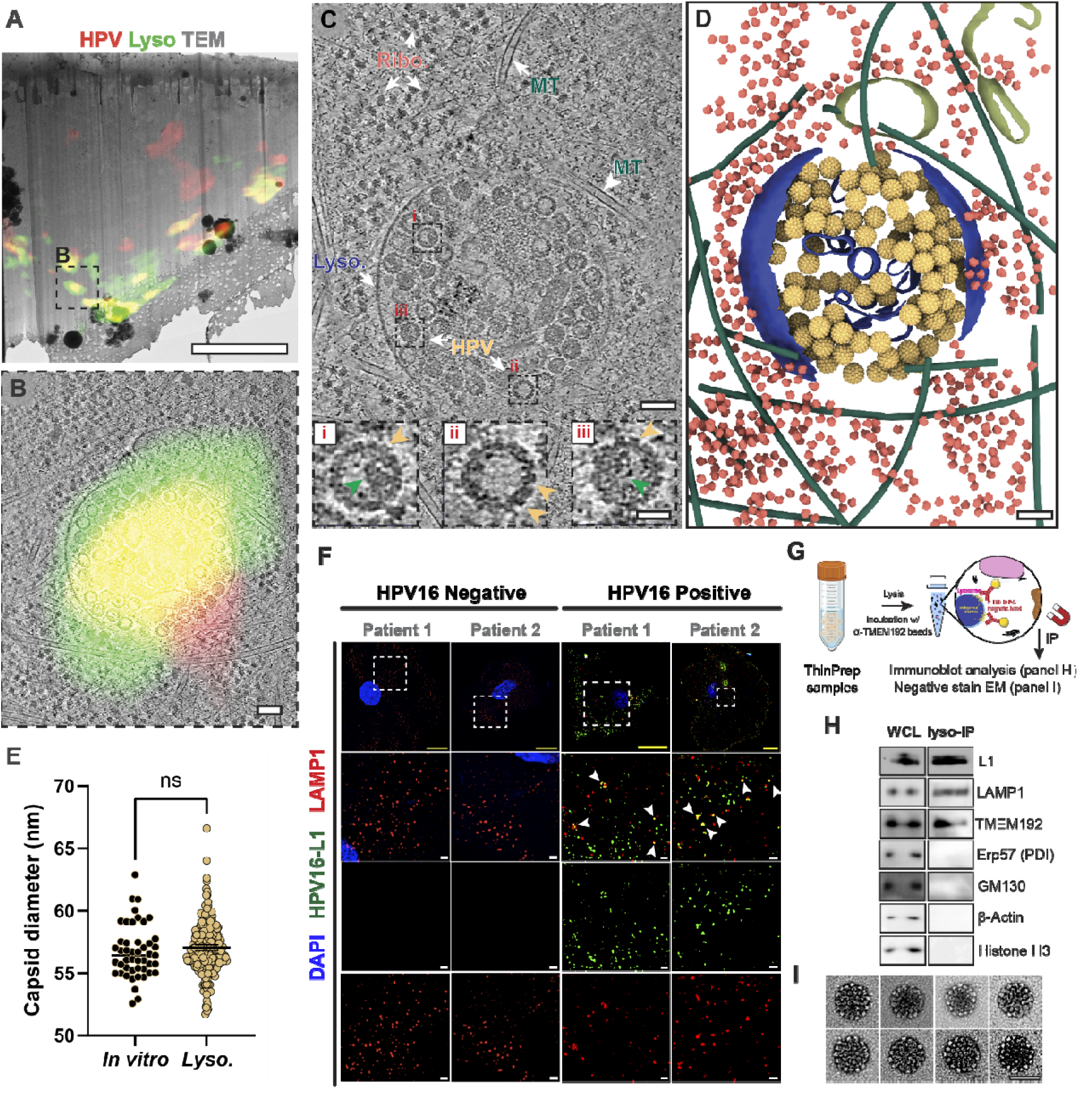
Human papillomavirus retains its morphological integrity within host lysosomes. **(A)** Cryogenic-correlative light and electron tomography (cryo-CLEM) analysis of HeLa cells infected with human papillomavirus (AF647-HPV, MOI=100). Overlay of the cryo-transmission electron microscopy (cryo-TEM, 6500×) and maximum intensity projection (MIP) cryo-confocal fluorescence image of a lamella through a HeLa cell, 16 hours post-infection (h.p.i.). Cryo-TEM (gray), and fluorescence signals from LysoTracker DND-26, and HPV, pseudocolored in green and red, respectively, are shown. The black dashed box highlights the region selected for tilt series acquisition. Scale bar: 5 µm. **(B)** Slice through a tomogram of the area highlighted in (A), overlaid with MIP cryo-confocal fluorescence signals: AF647-HPV (red) and LysoTracker (green). Scale bar: 100 nm. **(C)** Slice through a tomogram of the area highlighted in (A), highlighting intact cellular interiors of an HPV-infected HeLa cell. Cellular components such as ribosomes (Ribo), lysosomes (Lyso), microtubules (MT), and the virus (HPV) are clearly discernible and indicated by white arrows. Zoom-in views (5×, i-iii) highlight three HPV particles within the lysosomes. The distinct crown-like shape of the HPV L1 pentamer and the density of plasmid DNA are marked by yellow and green arrowheads, respectively. Scale bar: 25 nm, inset: 100 nm. See also corresponding Video S1. **(D)** 3D Segmentation of tomogram in (C) highlights HPV (yellow) and cellular features rendered as: lysosome membrane (dark blue), endoplasmic reticulum (ER, olive green), ribosome (salmon), microtubule (dark green). EMDB-5932 was used solely as a reference for visualizing HPV particles and does not imply that all particles are structurally identical or have uniformly packaged genomes. Scale bar: 100 nm. **(E)** Quantification of HPV capsid diameters. Diameter is defined as the maximum inter-pentamer (L1-L1) distance on the opposite side of the HPV capsid. Lysosomal HPV (n=288, 57.05 ± 1.96 nm) is similar in size to HPV purified *in vitro* (n=47, 56.71 ± 2.13 nm). (mean ± SD, Mann-Whitney t-test). **(F)** Representative confocal fluorescence microscopy images of ThinPrep samples (from patients 1 and 2). Healthy patients are on the left, while HPV16-infected patients are on the right. The top row shows low-magnification overviews, and the second row shows magnified views of the boxed regions, highlighting colocalization between HPV16 L1 and the lysosomal marker LAMP1. The third and fourth rows show the individual HPV16 L1 and LAMP1 channels, respectively. White arrowheads mark colocalization of L1 and LAMP1. Images are pseudo-colored as follows: Nucleus (DAPI, blue), HPV (L1 capsid protein, green), and lysosomes (LAMP1, magenta). Scale bars: yellow = 10 μm and white = 1 μm. **(G)** Schematic of tagless lysosome immunoprecipitation (lyso-IP) from ThinPrep samples obtained from HPV16-positive patients using TMEM192 antibody–coupled magnetic beads. **(H)** Representative immunoblot of whole-cell lysate (WCL, left) and tagless lyso-IP from HPV16-positive patient ThinPrep samples, probed for the HPV capsid protein L1 and the indicated organelle markers: LAMP1 and TMEM192 (lysosomes), Erp57 (endoplasmic reticulum), GM130 (Golgi), β-actin (cytosol), and histone H3 (nucleus). **(I)** Representative negative-stain electron micrographs of intact HPV particles isolated from lysosomal fractions of HPV16-positive patient ThinPrep samples. Scale bar: 50 nm.

Our observation that HPV virions remain intact within lysosomes suggests they may persist in infected patients for extended periods without clearance. To investigate this, we performed immunofluorescence staining on ThinPrep patient samples (see *Materials and Methods*). In samples from HPV16-positive patients, HPV16 L1 colocalized with the lysosomal marker LAMP1, indicating their presence within lysosomes, whereas no comparable signal was detected in healthy samples (**Figure 1F**). To assess the structural state of patient-derived lysosomal HPV, we immunoprecipitated lysosomes from ThinPrep samples of HPV16-positive patients using the tagless lyso-IP approach^86^ (**Figure 1G**). Immunoblot analysis confirmed enrichment of the lysosomal fraction, free of contamination from other organelles, and detected HPV16 L1 (**Figure 1H**). Subsequently, we performed negative-stain electron microscopy of these lysosomal fractions, which revealed several intact capsids (**Figure 1I**) resembling those observed in lysosomes from infected HeLa cells (**Figure 1C**). These findings suggest that contrary to the widely adopted model, HPV persists in the lysosome for an extended period and is unlikely to be completely degraded by lysosomal proteases.

LysoTracker is a weak base that becomes protonated and fluoresces in acidic environments ^87^. Our ability to precisely detect lysosomes in HPV-infected HeLa cells *in situ* suggests that these lysosomes retain their acidic pH, and yet the HPV capsids remain relatively intact. Intrigued by this finding, we next assessed whether lysosome-localized HPV remains structurally intact and infection-competent. We used lyso-IP to isolate lysosomes from HPV-infected HeLa cells that contained an HA tag at the C-terminus of an integral lysosomal membrane protein, TMEM192 ^88–90^ (**Figure 2A**). We performed immunoblotting to confirm the absence of other organelles, including the endoplasmic reticulum (ER), Golgi apparatus, nucleus, early endosomes (EE), and late endosomes (LE), and detected lysosomal proteins LAMP1 and TMEM192 as expected. We also detected both HPV capsid proteins L1 and L2 (**Figure 2B**). Next, we sonicated the lysosomes in an acidic buffer (pH 4.5) to release lysosome-resident HPV and evaluated its infectivity. Remarkably, using GFP expression to assess infection, we found that the lysosome-resident HPV exhibited infectivity similar to purified HPV PsV (**Figure 2C and S2C**). Because lysosomal HPV is infectious, we reasoned that artificially permeabilizing the lysosomal membrane with L-leucyl-L-leucine O-methyl ester (LLOMe) ^91–94^ might allow it to escape the lysosome and reach the cytosol, thereby providing an opportunity for the virus to enter the nucleus and cause infection. Indeed, LLOMe treatment of HPV-infected cells significantly increased infection (**Figure 2D**), further supporting the idea that lysosomal HPV is infectious. To determine whether this increase is specifically due to lysosome-resident HPV, rather than virus undergoing canonical Golgi-dependent trafficking, we inhibited the retrograde pathway with NAV-2729. We found that addition of LLOMe could rescue the NAV-2729 infection defect (**Figure S2D**). This rescue nevertheless required γ-secretase activity, indicating that L2-dependent membrane translocation or an associated γ-secretase dependent step remains necessary (**Figure S2E**). Additionally, LLOMe treatment failed to restore infection after cell-cycle arrest with the Cdk1 inhibitor RO-3306 (**Figure S2F**), indicating that the enhanced HPV infection resulting from artificial release of the virus from the lysosome still requires cell-cycle progression.

**Figure 2.**
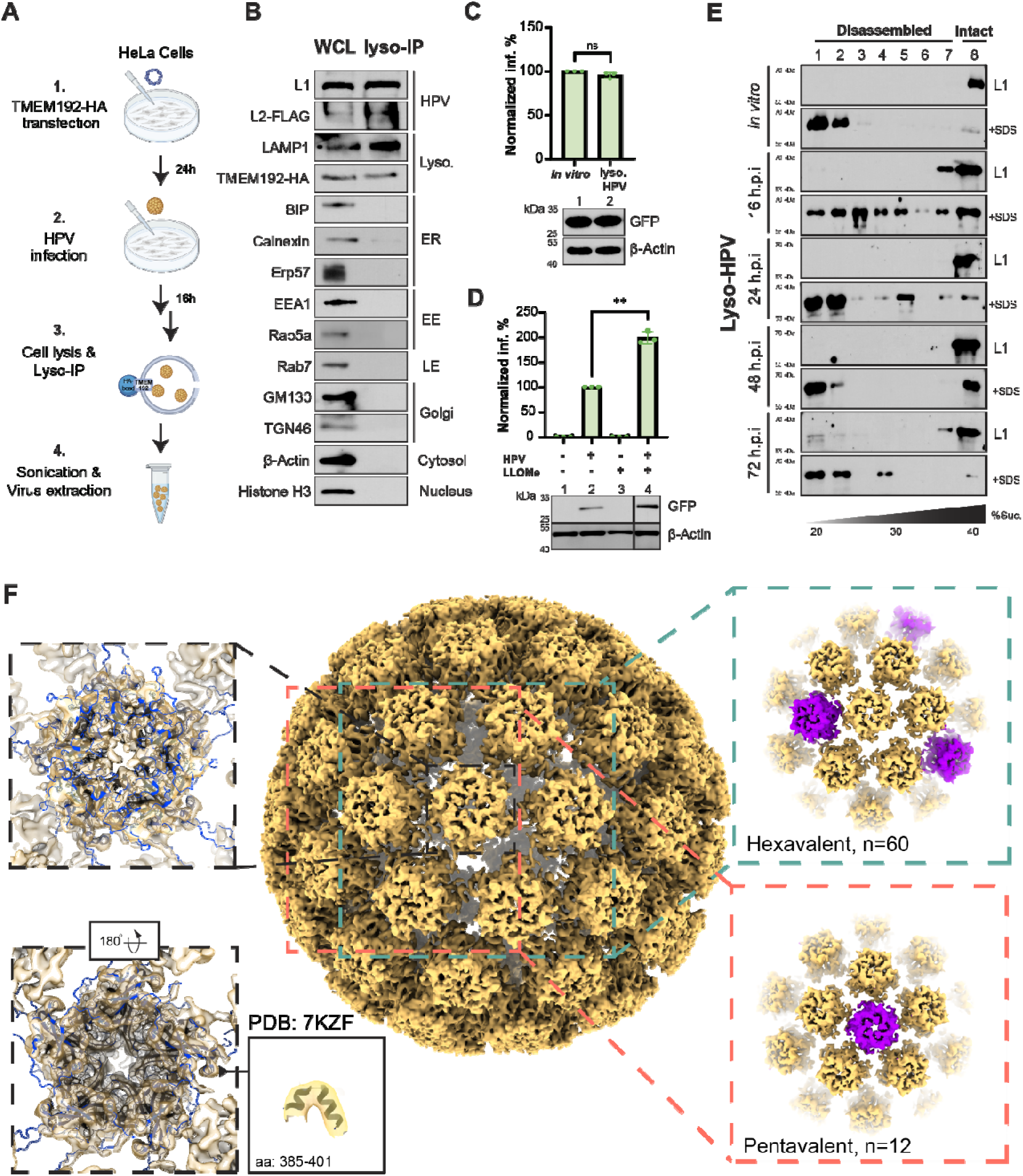
Lysosome-derived HPV is structurally intact. **(A)** Schematic of lysosomal immunoprecipitation (lyso-IP) workflow (see detailed explanation in Materials and Methods). Briefly, 1) HeLa cells are transfected with the TMEM192-HA plasmid. 2) 24 hours post-transfection (h.p.i.), cells are infected with HPV pseudovirus (MOI=50). 3) 16 hours later, cells are lysed and the lysate incubated with anti-HA beads for 5 minutes at room temperature. 4) Beads are incubated in a hypotonic pH 4.5 buffer for 20 minutes on ice and then briefly sonicated to rupture the lysosomes. **(B)** Representative immunoblot showing whole-cell lysate (WCL, left), immunoprecipitated beads (lyso-IP, right) against antibodies recognizing HPV capsid proteins (L1 and L2-FLAG) as well as different organelle markers, namely: LAMP1 and TMEM192-HA (lysosome), BIP, Calnexin, and Erp57 (ER), EEA1 and Rab5a (early endosome), Rab7 (late endosome) β-actin (cytosol), GM130 and TGN46 (Golgi), and Histone H3 (nucleus). **(C)** Representative immunoblot (bottom) and quantification (top) of reporter GFP expression in HPV-infected HeLa cells, prepared using standard procedures (*in vitro,* lane 1) or extracted from lysosomes (lyso. HPV, lane 2) (MOI =∼1). The amount of added virus was standardized based on L1 protein levels determined by immunoblotting (see Materials and Methods for details). At 48 h.p.i., cells were harvested, lysed, and the resulting whole-cell lysate was subjected to SDS-PAGE, followed by immunoblotting with antibodies recognizing GFP and β-actin as a loading control. GFP band intensities are normalized to β-actin in each sample. Total GFP band intensities per condition are normalized against the *in vitro sample* to give normalized infectivity (Normalized inf., %) (Welch’s t-test) (N = 3). **(D)** Immunoblots (bottom) and quantification (top) of GFP expression during HPV infection in the presence of LLOMe were performed as in (C). Cells were either treated with Dimethyl sulfoxide (DMSO, mock control, lane 1), HPV (lane 2), 1 mM of L-leucyl-L-leucine O-methyl ester (LLOMe, lane 3) or HPV and LLOMe (lane 4) (MOI=10). For quantification, DMSO was used as the normalization standard. (Welch’s t-test, ** p=0.0054) (N=3). **(E)** *In vitro* HPV or lysosome-extracted HPV (MOI=50) collected at the indicated time points from TMEM192-FLAG–expressing HeLa cells pretreated with 9 µM RO-3306 was layered onto a discontinuous 20%–40% sucrose gradient and subjected to centrifugation. Fractions were collected from the top of the gradient, and L1 distribution was assessed by immunoblotting. L1 detected in fractions 1–7 represents disassembled viral material, whereas L1 in fraction 8 corresponds to intact virions. (**F**) Single particle cryo-electron microscopy analysis of lysosomal-extracted HPV. Reconstruction from 8,561 particles yielded a 6.3 Å Coulomb potential map (center, yellow) showing the characteristic icosahedral architecture of HPV. L1 capsomer forms both hexavalent (inset right, top) and pentavalent (inset right, bottom) interactions with other L1 capsomers. The pentavalent L1 capsomer is highlighted in magenta. A rigid-body fit of the L1 structure (PDB: 7KZF) into the L1 capsomer of the lysosomal HPV map (inset, top left) is shown with a view from within the virus to the outside. Additionally, a close-up view of the selected regions (aa:385-401) of the capsomer density is shown in the inset (bottom left).

To evaluate capsid integrity across the lysosome-associated population, we applied a previously established sucrose cushion sedimentation assay ^43,95,96^. Briefly, we layered lysosome-derived HPV onto a discontinuous 20%–40% sucrose gradient and subjected it to ultracentrifugation. Under these conditions, disassembled viral material would remain in the upper fractions (fractions 1–7), whereas intact particles sedimented to the bottom fraction (fraction 8). To validate the assay’s ability to distinguish intact from disassembled viral capsids, we used *in vitro* HPV. We detected this exclusively in fraction 8, consistent with an intact capsid; upon SDS treatment, L1 shifted into the lighter fractions (**Figure 2E**, top panel). When we analyzed lysosome-associated HPV recovered at 16, 24, 48, and 72 h.p.i., most L1 remained in fraction 8 at every time point examined (**Figure 2E**). We obtained similar results in HEK293 and HaCaT cells (**Figures S2G and S2H**, respectively), indicating that this phenotype is not restricted to a single cell type. Together, these findings show that lysosomal HPV capsids remain intact for at least 48 h, with capsid disassembly beginning at 72 h, the latest time point analyzed.

To further substantiate this conclusion, we employed single-particle cryo-electron microscopy (cryo-EM) to determine the structure of lysosomal HPV. Cryo-EM analysis revealed that the lysosomal HPV consists of 72 capsomers, each composed of 5 L1 proteins, similar to the previously determined structure of HPV ^21,22^. We found that 12 capsomers form pentavalent interactions with neighboring capsomers (magenta, **Figure 2F**), while 60 L1 capsomers form hexavalent interactions. The previously determined capsomer structure (PDB: 7KZF) fits very well in our Coulomb potential map (**Figure 2F**, insets to the left). Our map aligns perfectly well with the previously determined cryo-EM structure of the isolated HPV capsid (**Figure S4D**, EMD-23081). Therefore, at 6.3 Å, our reconstruction of lysosomal HPV provided sufficient architectural detail for this study, and we chose not to pursue further data collection or resolution improvement.

Collectively, our results demonstrate that HPV virions in the lysosome maintain their structural integrity and may not undergo significant disassembly or degradation, as previously proposed ^31–44^. Additionally, our data support the presence of structurally intact and infectious virions in lysosomes under the conditions tested, and these findings should motivate further investigation into the state and fate of the viral capsid within the presumed non-productive trafficking pathways such as lysosomes.

### Intact HPV traffics to the Golgi, where it buds into COPI-coated transport vesicles

The prevailing paradigm in HPV trafficking and infection is that the L1 capsid progressively disassembles as the virus undergoes retrograde transport to reach the TGN/Golgi ^51,67–70^. L1 capsid disassembly is thought to expose the hidden L2 capsid, allowing it to insert into and translocate across the host membrane, a process necessary to guide the virus toward productive infection ^46,48,97,98^. We asked whether intact HPV capsids are trafficked to the TGN/Golgi. To visualize HPV architecture during retrograde trafficking, we analyzed tomograms of HeLa cells infected with HPV at 22 h.p.i., specifically at sites of colocalization between AF647-HPV and HA-COPγ1-mCherry fluorescence. Strikingly, we observed intact HPV capsids in the Golgi lumen (**Figures 3A-J and S3I; Videos S2 and S3**). Detailed examination of various tomographic Z-slices through HPV particles revealed apparent individual L1 pentamers and captured particles in the process of budding, revealing the formation of the COPI coat at the initiation of budding from the Golgi membrane (**Figures 3C-E and 3H-J**; a schematic of the Golgi membrane, COPI coat, and HPV is shown on the right). We also observed a fully budded vesicle harboring HPV, surrounded by a partial COPI coat, localized adjacent to the discernible Golgi stacks (**Figures 3K, 3L, and S3I**). Finally, across several tomograms, we captured snapshots of HPV in the cellular milieu, thereby allowing us to pseudo-reconstruct the entire budding process. We categorized the process into four distinct steps: HPV capsids initially reside in the Golgi lumen, followed by COPI coat recruitment and membrane budding, which leads to the formation of a COPI-coated vesicle containing the virus (**Figure 3M**). Eventually, the COPI coat disassembles from the budded vesicles (**Figure 3M**, right).

**Figure 3.**
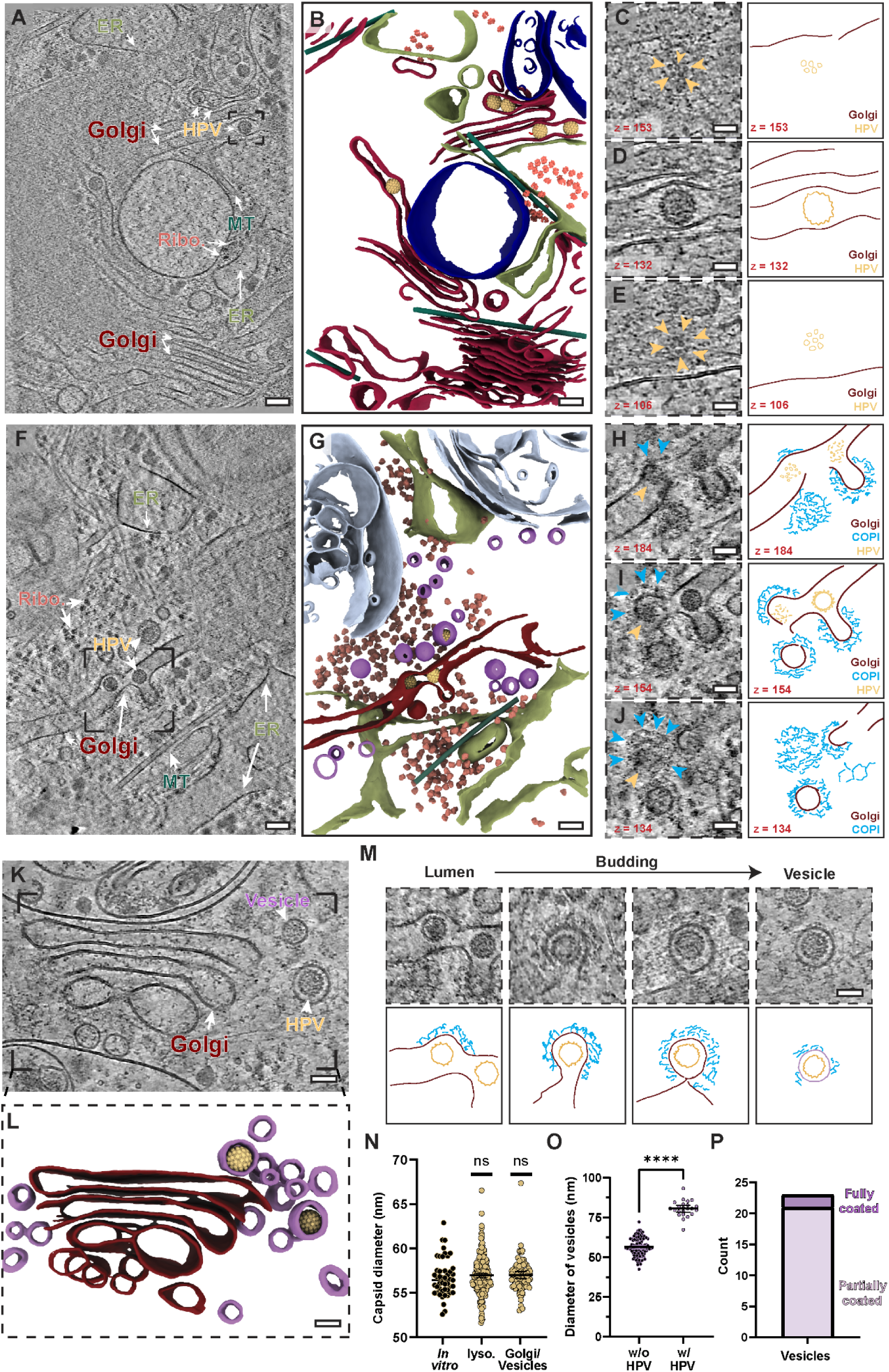
*In situ* structural analysis of HPV at the Golgi. **(A, F)** Slice through a reconstructed and NAD-filtered tomogram collected at the site of COPγ1 and HPV colocalization (see Figure S3), revealing the cellular milieu at the Golgi (MOI =100). HPV particles are present within the Golgi lumen. HPV (yellow) and cellular components such as the endoplasmic reticulum (ER), Golgi, microtubules (MT), and ribosomes (Ribo.) are depicted by white arrows. Scale bar: 100 nm. See also corresponding Videos S2 (panel F) and S3 (panel A). **(B, G)** 3D segmentation of the tomogram in (A) showing HPV (yellow) and key cellular features rendered as: Golgi (maroon), Multivesicular bodies (MVB, light blue), lysosome membrane (dark blue), ER (olive green), vesicle (purple), ribosome (salmon), and microtubules (dark green). Scale bar: 100 nm. **(C-E)** Left: close-up views of the boxed region in (A), showing three different tomographic slices at 153 (C), 132 (D), and 106 (E), showing an individual HPV L1 pentamer (yellow arrowheads). (D) shows the slice containing HPV in the Golgi stack. (C) and (E) show slices approximately 22.3 nm above and 27.6 nm below the plane in (D), respectively. Right: Cartoon schematic of the Golgi (maroon) and HPV (yellow) corresponding to the slices on the left. Scale bar: 50 nm. **(H-J)** Left: close-up views of the boxed region in (F), showing three different tomographic slices at 184 (H), 154 (I), and 134 (J), showing the formation of the COPI coat (blue arrowheads) around the HPV particles (yellow arrowheads) ready to exit the Golgi lumen. (I) shows the slice containing HPV in the Golgi lumen and an apparent COPI coat on the membrane. (H) and (J) show slices approximately 31.9 nm above and 21.2 nm below the plane in (I), respectively. Right: Cartoon schematic of the COPI coat (cyan), Golgi (maroon), and HPV (yellow) corresponding to the slices on the left. Scale bar: 50 nm. **(K)** Slice through a tomogram showing a characteristic Golgi stack and the HPV particles post exit from the Golgi, within vesicles. Scale bar: 50 nm. (**L**) 3D segmentation of the tomogram in (K), highlighting the Golgi (maroon) and the vesicles (lilac) harboring HPV particles. Scale bar: 50 nm. (**M**) Top: representative snapshots depicting HPV exiting the Golgi. Slices through different tomograms, from left to right, show formation of the COPI coat, expansion of the coat, a fully mature vesicle about to be scissioned from the Golgi, and HPV in a vesicle derived from the Golgi. Scale bar: 50 nm. Bottom: Cartoon schematic of the various stages of HPV exit from the Golgi. COPI (cyan), Golgi (maroon), and HPV (yellow) are shown. **(N)** Quantification of the capsid diameters of purified HPV (left, black circles), lysosomal HPV (middle, yellow circles), and HPV within the Golgi or coated vesicles (yellow circles, right). Diameter is defined as the maximum inter-pentamer (L1-L1) distance on the opposite side of the HPV capsid. HPV within the Golgi and vesicles (n=86, 57.17 ± 1.85 nm) shows no change in diameter, in comparison to purified *in vitro* (n=47, 56.71 ± 2.13 nm) or lysosomal HPV (n=288, 57.05 ± 1.96). (mean ± SD, Mann-Whitney t-test) **(O)** Quantification of COPI vesicle diameters, measured as the maximum distance between the centers of opposing membrane leaflets, with HPV (n=23, 80.53 ± 5.11 nm) and without HPV (n=104, 56.40 ± 5.76 nm) (mean ± SD, Mann-Whitney t-test,**** p<0.0001) **(P)** Bar plot illustrating the number of observed instances of vesicles with a full COPI coat (2/23), or partial coat (21/23) in cryo-ET data.

Throughout all stages of this process, HPV capsids remain morphologically assembled, as defined by the observed crowns of L1 pentamers in the tomographic slices. We measured HPV particle diameters in the Golgi compartment and COPI-coated vesicles. We found no difference between the mean capsid diameter in the Golgi/vesicle populations (57.17 ± 1.85 nm), lysosomal HPV (57.05 ± 1.96 nm), and purified HPV particles (56.71 ± 2.13 nm) (**Figure 3N**), revealing that the virus capsids can remain structurally intact without the need to undergo extensive disassembly during trafficking to the Golgi.

We next assessed the diameter of COPI-coated vesicles with and without the HPV cargo. Vesicles lacking virus cargo had a mean diameter of 56.40 ± 5.76 nm (**Figure 3O**), consistent with previous reports ^99–101^. By contrast, COPI-coated vesicles containing HPV were significantly larger, with a mean diameter of 80.53 ± 5.11 nm. This substantial increase in size shows that COPI vesicles can adapt to accommodate unusually large cargo. Finally, we also examined the state of COPI-coated vesicles containing HPV. We found that 21 out of 23 vesicles exhibited partial disassembly of the COPI coat, while only two remained fully coated (**Figure 3P**). This finding is consistent with the established maturation mechanism of COPI-coated vesicles in which the COPI coat dissociates from the membrane following vesicle formation ^102–105^. Taken together, we unequivocally show that intact HPV capsids bud from the Golgi in COPI-coated vesicles.

### Proteinaceous structural bridges link HPV to host membranes during retrograde trafficking

We next asked whether Golgi- or vesicle-resident HPV has distinct hallmarks (**Figure 4**). We consistently observed distinct protein densities spanning the Golgi lumen connecting the HPV and the Golgi membranes [representative examples of three independent tomograms containing Golgi-resident HPV with bridging densities highlighted in **Figure 4C, 4H, and 4M**, red arrowheads; and corresponding segmentations in **Figure 4D, 4E, 4I, 4J, 4N, and 4O]**. Each instance showed multiple discrete densities bridging the virus capsid and the overlaying membrane surface. Similarly, we also found connector densities between HPV and the encapsulating membranes in fully budded vesicles (**Figure 4P**). Again, the tomograms revealed comparable proteinaceous densities at the interface between the virus capsid and the vesicular membrane, with several apparent contact points. We quantified both the length and width of these bridging densities, in which the length is defined as the distance from the surface of the virus to the inner leaflet of the membrane, and the width as the span from one end of the proteinaceous density to the other. On average, the Golgi bridging densities measured 8.58 ± 2.41 nm in length and 2.41 ± 0.46 nm in width (**Figure 4Q**), while the vesicle bridging densities measured 7.27 ± 1.21 nm in length and 2.33 ± 0.51 nm in width (**Figure 4R).** Given their relatively consistent length, we next asked whether these densities constrain HPV particles to an off-center position within the larger Golgi-derived vesicles. We therefore performed a quantitative centroid analysis of HPV particles and their enclosing vesicles in our tomograms, defining viral offset as the distance between the particle center and the vesicle center, as illustrated in the schematic. Our analysis revealed that the virus’s center is systematically offset from the vesicle’s center, indicating an intrinsic asymmetry in virus–vesicle interactions that skews the virus towards one side of the vesicle (**Figure 4S**).

**Figure 4.**
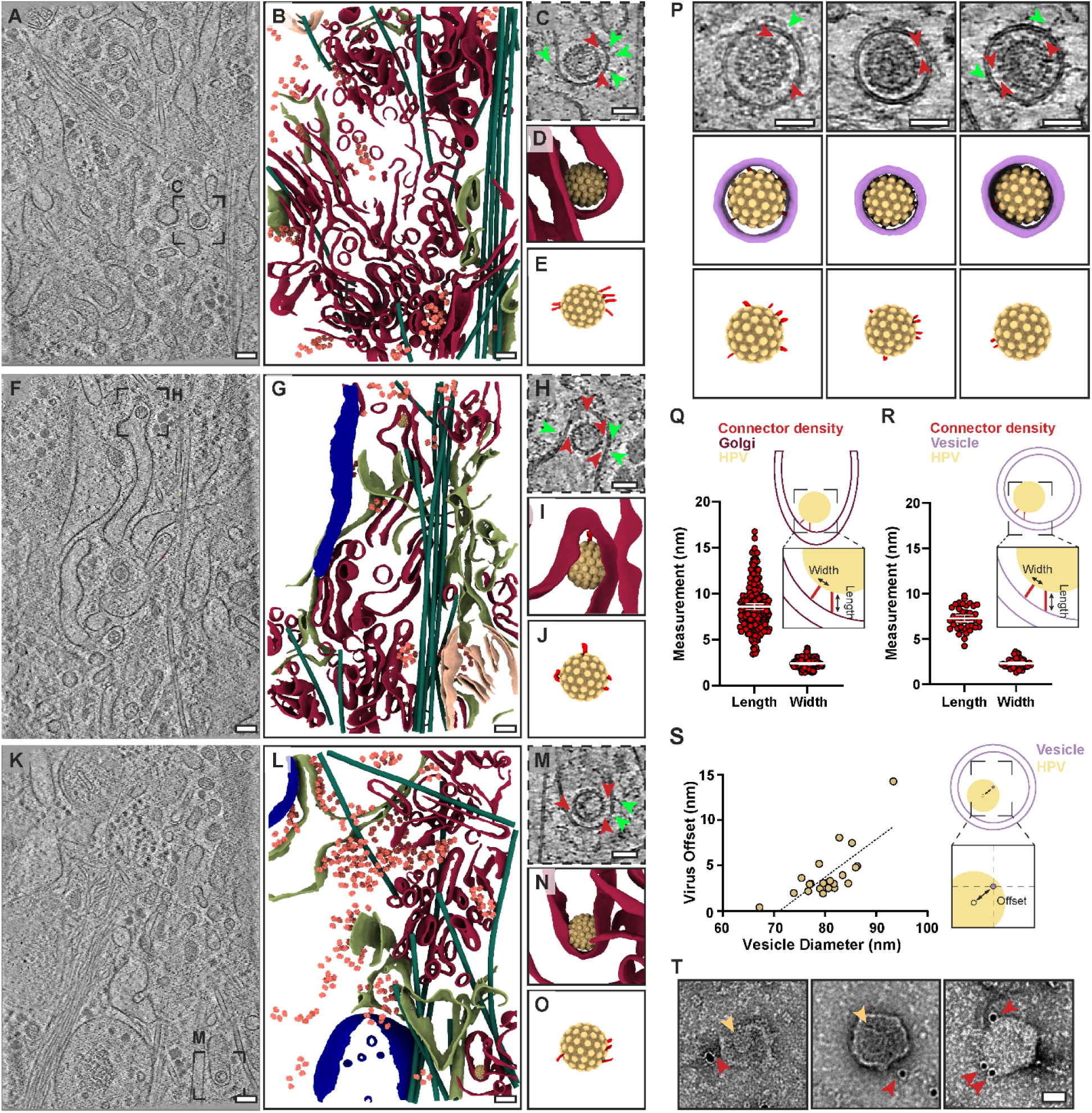
Protein densities bridge the viral capsid to Golgi and COPI-coated vesicle membranes. **(A, F, K)** Slices through three independently reconstructed and NAD-filtered tomograms showing the cellular environment surrounding the Golgi (MOI=100). HPV particles are visible within the Golgi lumen. Scale bars, 100 nm. **(B, G, L)** 3D segmentations of tomograms shown in (A), (F), (K), respectively. Cellular features and viral particles are rendered as: Golgi (maroon), lysosome membrane (dark blue), ER (olive green), mitochondria (peach), ribosome (salmon), microtubules (dark green), and HPV (yellow). Scale bar: 100 nm. **(C, H, M)** Magnified views of the boxed region in (A), (F), and (K), respectively, showing multiple proteinaceous densities (red arrowheads) bridging the viral capsid and the Golgi membrane. Bright green arrowheads indicate extension of these bridging densities outside the Golgi lumen. **(D, E, I, J, N, O)** Corresponding 3D segmentations are shown with the membrane (D, I, N) or without the membrane (E, J, O) of (C), (H), (M), respectively. The cellular features and viral particles are rendered as: Golgi (maroon), HPV (yellow), and proteinaceous density (red). Scale bars: 50 nm. **(P)** Top, representative slices through reconstructed and NAD-filtered tomograms showing HPV particles within COPI-coated vesicles (MOI=100). Red arrowheads indicate multiple proteinaceous densities bridging the viral capsid and vesicle membrane, and bright green arrowheads indicate extensions of these densities. Middle and bottom row: the corresponding 3D segmentation shown with and without membrane, highlighting cellular features and viral particles rendered as: vesicle (purple), HPV (yellow), and proteinaceous density (red). Scale bars: 50 nm. **(Q, R)** Schematics and quantification of the dimensions of bridging densities observed within the Golgi lumen (Q) or vesicles (R). Golgi-associated densities measured 8.58 ± 2.41 nm in length and 2.41 ± 0.46 nm in width, whereas vesicle-associated densities measured 7.27 ± 1.21 nm in length and 2.33 ± 0.51 nm in width. Length was defined as the distance from the viral surface to the inner leaflet of the membrane, and width as the lateral span of the density. Data are presented as mean ± SD (Golgi, n = 236 densities; vesicles, n = 45 densities). **(S)** Schematic and scatter plot showing the positional offset of each viral particle relative to its diameter within the vesicle. Each point represents an individual HPV-containing vesicle (n = 23). **(T)** Representative negative stain electron micrographs of immunogold (anti-FLAG) labeled vesicles containing HPV, isolated via FLAG-IP. Red arrowheads indicate gold beads recognizing the C-terminus of HPV L2 protein and decorating the surface of the vesicle. The yellow arrowhead indicates an HPV particle located within the vesicle. Scale bar: 50 nm (MOI =50).

While debated, the current understanding is that during retrograde trafficking, the HPV L2 capsid is exposed from within the HPV L1 capsid, enabling it to insert into and translocate across host membranes to recruit cytosolic factors that guide viral transport ^45,46,48,97^. In some tomogram slices, we observe the connector density extending across the vesicle membrane bilayer and into the cytoplasm (**Figure 4C, 4H, 4M, and S3I,** light green arrowheads). Additionally, while up to 72 copies of L2 have been estimated to be present per HPV capsid ^20–22^, we never observed more than 15 connector densities per viral particle in our *in situ* analysis. Structural predictions show that HPV L2 is mostly disordered, with the central structural region (CSR) of 109 amino acids located between the disordered N- and C-termini, with approximate dimensions of 6.2 nm and 2.8-4 nm along the long and short axes, respectively (**Figure S5**). These dimensions align reasonably with those of the proteinaceous bridges (**Figure 4Q, R**).

We therefore surmised that the CSR of the L2 capsid protein may form this structural bridge between the virus and the encapsulating vesicle and may orient in a way that the distal L2 C-terminus would be accessible on the exterior of the vesicle (**Figure S5F**). To assess whether the C-terminus of L2 is exposed on the exterior of the membrane, we took advantage of the 3×- FLAG epitope engineered at the C-terminus of L2 in our HPV PsV. We used FLAG immunoprecipitation to rapidly isolate intact HPV-containing vesicles. Following isolation, we probed for the accessibility of the L2 C-terminus on the vesicle surface using gold-labeled anti-FLAG M2 antibody. We observed gold beads decorating the exterior of isolated vesicles (**Figure 4T**, red arrowheads). These results demonstrate that the C-terminus of L2 is exposed on the exterior of the HPV-containing vesicles. Collectively, these findings highlight a direct physical connection between HPV and host vesicular compartments which could possibly be mediated by HPV L2.

## Discussion

Globally, HPV is responsible for 4.5% of all human cancers and remains a significant public health threat ^7,8^. While highly effective prophylactic HPV vaccines exist, the majority of individuals worldwide have not been vaccinated; current estimates suggest that 13 million new infections are reported annually in the U.S. alone, and the average lifetime probability of acquiring HPV among those with at least one opposite-sex partner is 84.6% for women and 91.3% for men ^14,15,106,107^. Several reports also suggest that older people are more susceptible to HPV reactivation ^108–115^. Clarifying the cellular routes of HPV infection will therefore provide better insights into the design of new anti-HPV strategies. Here, using *in situ* cryo-ET, we provide additional key molecular insights into decisive events in infection.

### HPV at the lysosome

After receptor-mediated endocytosis, HPV reaches the endosome, where L1 capsids are thought to undergo partial disassembly ^31–44^. Several host factors, including specific Rab GTPases, the ESCRT machinery (i.e., Tsg101 and Vps4), and the V-ATPase, have been implicated in promoting viral capsid disassembly in the endosome ^36,40,37,35,39,41^. However, whether these factors directly promote capsid disassembly or act indirectly by maintaining endosomal homeostasis remains unclear. An alternative model proposes that endosomal acidification induces capsid “loosening,” thereby initiating disassembly. This idea is primarily based on *in vitro* studies demonstrating that low pH increases the accessibility of the encapsulated viral genome to DNase I ^32^. Support for this model, however, comes predominantly from pseudovirions carrying reporter plasmids, whose genome composition and packaging density differ from those of native or quasiviral HPV genomes; because these properties can influence capsid stability, the two preparations may not respond equivalently to acidification. Beyond the endosome, HPV can also be delivered to the lysosome, where it is thought to be disassembled and degraded^30,32–34,38,65,66^.

Here, we demonstrate conclusively, in both cell culture models and samples from living patients, that lysosome-localized HPV can remain morphologically intact.

We show that the average diameter of lysosomal HPV is indistinguishable from that of native or Golgi-localized virus (**Figure 3N**). By isolating lysosomal HPV at pH 4.5 and determining its cryo-EM structure, we show that the capsid withstands the acidic lysosomal environment (**Figure 2F**). This finding agrees with our confocal microscopy data, which show high colocalization between L1 and the lysosomal marker LAMP1, even at 72 hours post-infection (**Figure S2A and B**). To further validate that this paradigm holds in human infection, we analyzed samples from patients positive for HPV16. We observed a fraction of HPV colocalizing with LAMP1 (**Figure 1F**), and lysosomes isolated from these samples revealed several intact capsids (**Figure 1I)**, indicating the virus’s resilience to lysosomal acidity and proteases. Collectively, these findings not only strongly support HeLa cells as a robust model for HPV research but also show that HPV is far more resistant to lysosomal degradation than previously appreciated. In line with this, previous cryo-EM studies of purified HPV, incubated in buffers spanning near-neutral to moderately acidic conditions (pH 7.11, 6.08, 5.45) showed no detectable change in capsid diameter, with pronounced capsid degradation only occurring at pH 2 ^116^. Finally, we also showed that the lysosomal pool of HPV is as infective as the purified pseudovirus (**Figure 2C and S2C**). Our findings reveal that lysosomes do not degrade all internalized HPV and suggest that they may serve as a reservoir of infection-competent particles. However, whether these particles normally exit lysosomes and re-enter a productive infection pathway remains to be determined.

What might be the advantage of storing HPV in the lysosome? One possibility is that, should the endosome be overcrowded with HPV, which prevents its efficient delivery to the TGN/Golgi for productive infection, endosome-localized HPV might instead be directed to the lysosome for temporary storage and only be re-routed back to the endosome when conditions are less crowded. In this manner, there would be a sustained supply of HPV particles that can be used for infection. This proposed lysosome-endosome-TGN/Golgi transport pathway has precedent. For instance, in yeast, the Atg27 membrane protein is recycled from the vacuole to the endosome and then targeted to the Golgi by the retromer ^117^. The possible involvement of the lysosome during HPV entry is not exclusive to this virus. For instance, coronaviruses, including SARS-CoV-2, use the lysosome for unconventional egress by exploiting the harsh conditions within this compartment ^118^. In contrast, the Ebola virus glycoprotein is cleaved by lysosomal proteases, allowing the virus and lysosomal membranes to fuse and deliver its genome into the cytosol ^119^.

### HPV at the Golgi

Regardless of the precise mechanism, disassembly is thought to expose the L2 capsid protein, which subsequently inserts across the endosomal membrane and becomes accessible to the cytosol ^45,46,48,97^. The cytosol-exposed region of L2, in turn, recruits trafficking factors that guide the virus to the TGN, committing it to the productive infection route ^50–58^. We previously demonstrated that at the TGN, HPV L2 binds directly to the COPI coat complex, enabling the virus to undergo retrograde transport to and across the Golgi stacks before reaching the nucleus to cause infection ^60^. However, how COPI promotes retrograde trafficking of HPV is not entirely clear. Importantly, in this scenario, HPV is expected to be at least partially disassembled upon arrival at the Golgi.

Our study demonstrates that the HPV capsid retains its structural integrity in the Golgi lumen, on the COPI-coated Golgi membrane, during budding initiation, and in Golgi-derived budded vesicles (**Figure 3**). In these steps, quantitative measurements reveal no significant differences in HPV capsid diameters relative to those of native virions, indicating that the capsid architecture remains largely unchanged throughout retrograde transport from the cell surface to the Golgi (**Figure 3N)**.

Serial snapshots of intact HPV in the Golgi lumen, COPI-coated pits in the Golgi membrane, and fully-budded vesicles (**Figure 3M**) strongly suggest that the COPI budding machinery is responsible for generating membrane-bound vesicles harboring HPV that are used for retrograde trafficking. Our *in situ* cryo-ET data not only provide visual evidence but also validate our previous work, suggesting that HPV L2 binds to COPI during retrograde trafficking across the Golgi, thereby facilitating infection^60^. *In situ* structural analysis of COPI coats in *Chlamydomonas reinhardtii* indicated the presence of cargo bound beneath the B’ subunit of COPI ^100^; nevertheless, the precise nature of cargo remained unknown. Here, we provide the first direct *in situ* evidence of a virus exploiting the mammalian COPI machinery for retrograde transport (**Figure 3**). The only previous study demonstrating *bona fide* cargo within coated vesicles by *in situ* cryo-ET was of COPII-coated vesicles; however, it used a reconstituted approach with purified proteins and cell-derived membranes rather than native cellular conditions ^120^.

Our cryo-ET analysis revealed that HPV-containing vesicles were significantly larger than those without viral cargo. While vesicles devoid of HPV are approximately 56 nm in diameter and are consistent with previous reports ^99–101^, HPV-containing vesicles expand to approximately 80.5 nm in diameter (**Figure 3O**). To our knowledge, this is the first report showing that mammalian COPI-coated vesicles can adapt in size, growing to accommodate the cargo. This ability likely reflects the inherent structural flexibility of the COPI coatomer complex, which exists in the cytosol as a dynamic, highly flexible assembled protein complex and is recruited to membranes as a heteroheptameric unit during vesicle formation ^99,100,121–125^. The proposed flexibility in COPI-derived vesicles is reminiscent of that in COPII-coated vesicles, which can also expand to accommodate large cargoes such as procollagen ^126–128^.

Furthermore, we identified distinct proteinaceous densities that bridge HPV particles to adjacent host membranes within the Golgi lumen and budded vesicles (**Figure 4**). While the precise molecular identity of these densities remains to be established, we show that HPV engages in asymmetric interactions with the Golgi and vesicle membranes via this density. One possibility is that the density comprises capsid protein L2, which undergoes cleavage ^21,24–28^ during entry and protrudes across endosomal membranes to recruit cytosolic trafficking factors. Consistent with this interpretation, we did not observe such bridging densities in COPI-coated vesicles lacking HPV, and the dimensions approximate those of the L2 central structured region (CSR), as predicted by AlphaFold 3 (**Figure S5C**). However, because a high-resolution structure of L2 in this context is unavailable, we cannot confidently assign these densities to L2 or to a specific L2 region. Interestingly, although the proteinaceous bridging densities in the Golgi and vesicles had comparable average widths, the Golgi-associated densities were longer on average. This difference may reflect conformational remodeling of the bridging complex during vesicle budding, although its mechanistic significance remains to be determined.

A recent report showed that different disordered L2 segments can form finger-like projections that extend into the central pore of an L1 pentamer; in some models, the L2 C-terminus traverses this pore and becomes exposed on the capsid surface. Consistent with this model, the authors detected an average of seven surface-exposed L2 C-termini per capsid. Together, these findings support a mechanism in which L2 protrudes through intact L1 pentamers without requiring extensive capsid disassembly^129^. Thus, our cryo-ET images may capture different states of HPV L2. Alternatively, the bridging densities may represent host factors recruited in response to HPV presence or complexes comprising L2 and host proteins. Different L2 molecules within the same particle could also engage distinct host factors at successive stages of trafficking. Defining the molecular composition of these densities and the temporal regulation of their interactions will require further investigation.

Whether HPV is disassembled after exiting the Golgi *en route* to the nucleus is unclear and deserves future investigation. In this context, a previous study using standard TEM showed that HPV particles within a membrane-bound vesicle can be found in the nucleus ^130^. However, the lower resolution of this approach cannot determine if the virus is fully intact. Moreover, a recent study showed that nuclear phospholipases promote release of the HPV genome from transport vesicles, supporting a model in which genome release and potentially late-stage capsid disassembly occur in the nucleus, downstream of Golgi exit^131^.

In sum, we report the first high-resolution molecular snapshots of HPV trafficking in its native cellular environment. Although current models suggest limited capsid disassembly, our results show that HPV capsids can remain largely intact during retrograde trafficking to the Golgi. Our data further support a coat-dependent budding mechanism for generating HPV-containing transport vesicles and demonstrate that the lysosome does not degrade HPV entirely but instead harbors a persistent pool of particles that retain infectivity following isolation. Although it remains unknown whether these particles normally exit lysosomes, our findings raise the possibility that lysosomes may serve as a storage compartment for infection-competent HPV. These new insights should guide future efforts to identify novel targets for anti-HPV intervention and support the IPVS Consensus that HPV can remain undetected after infection ^132^.

## Acknowledgements

We thank all members of the Tsai and Mosalaganti laboratories for their helpful suggestions and discussions. We thank the University of Michigan cryo-electron microscopy facility for their assistance in data collection workflows. We thank Dr. Vikas Navratna and Dr. Tai-Ting Woo for their help in refining the biochemistry protocols. We thank Ms. Stella Mulrenin for her contribution to pseudovirus production. Some schematics were created in BioRender by Sutanto, R. (2025).

## Funding

This work was supported by American Heart Association Grant 25PRE1361891 (R.S.); National Institutes of Health 5R01AI150897-05 (B.T.); National Institutes of Health 1DP2GM150019-01, and Klatskin Sutker Discovery Fund (S.M.); National Institutes of Health S10OD030275 and the Arnold and Mabel Beckman Foundation award to the University of Michigan Cryo-EM facility. The funders had no role in the study design, data collection, analysis, or the content and publication of this manuscript.

## Author contributions

R.S. performed most of the experiments, including preparing and analyzing immunoblots, Lyso-IP, FLAG-IP, negative staining, immunogold labeling experiments, immunofluorescence, confocal microscopy, and the cryo-CLEM/cryo-ET workflow, including cryo-FIB milling, cryo-ET tilt series collection, tilt series reconstruction, and segmentation. S.M. prepared cryo-EM samples, processed the single-particle cryo-EM dataset, performed the AlphaFold predictions, and prepared the movies. R.S. and S.M. analyzed the cryo-ET data. J.R. collected single-particle cryo-EM data. P.L. and D.M.H. collected and processed the ThinPrep patient samples. C.Y.C. performed immunofluorescence staining and imaging on ThinPrep patient samples. R.S. wrote the first draft of the manuscript. R.S and S.M. made the figures. R.S., D.M.H., B.T., and S.M. conceptualized the study, performed analysis, supervised the project, and finalized the manuscript.

## Competing interests

The authors declare that they have no competing interests.

## Notes

### Competing Interest Statement

The authors have declared no competing interest.

### Summary of Updates

The manuscript has been revised to include additional data supporting the claims.

