## Supplementary Information for "Redefining paradigms of cellular HPV trafficking *in situ*"

This file includes:

Figs. S1-S5

Materials and Methods

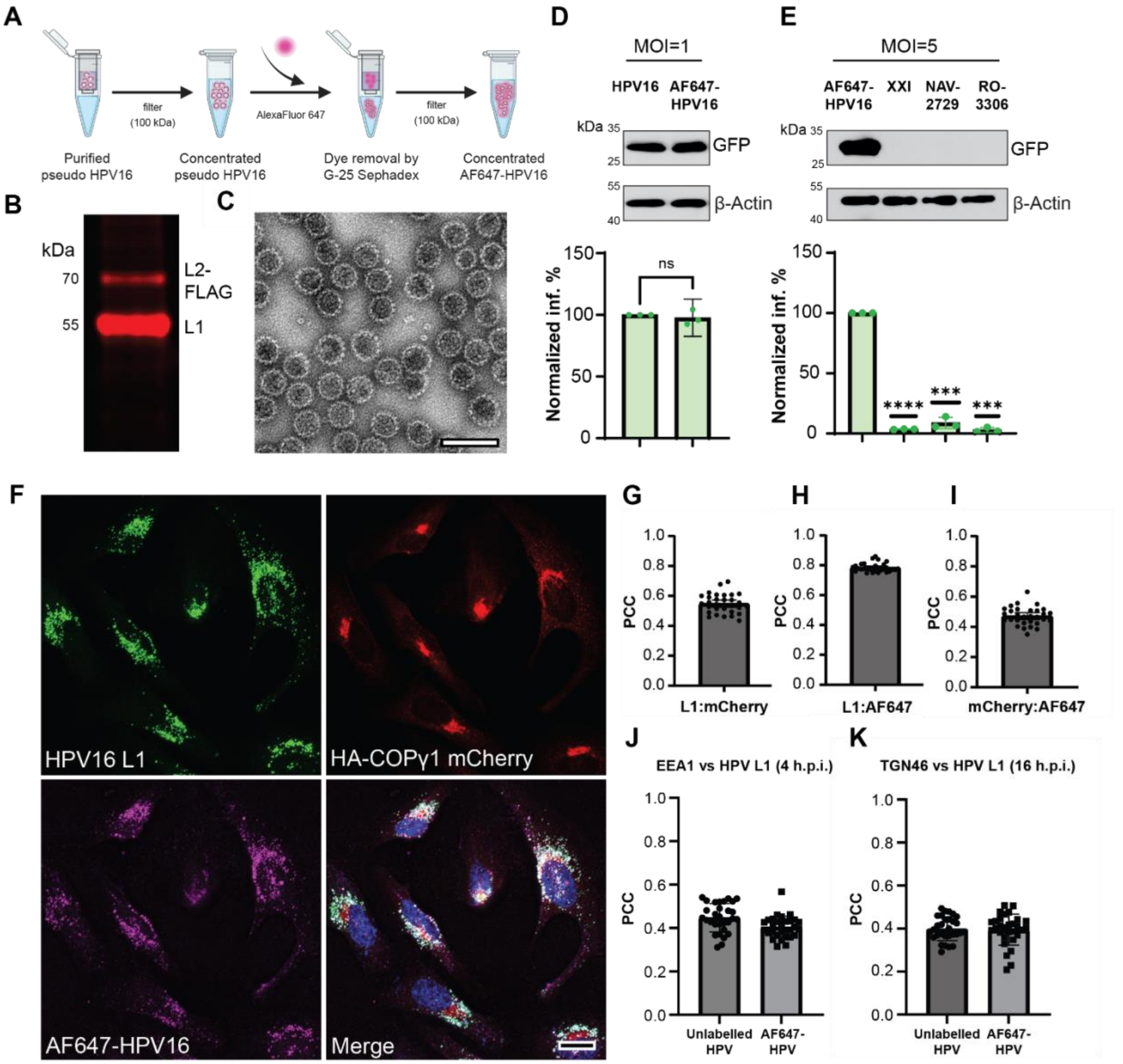


**Figure S1. Characterization of fluorescently labeled HPV. (A)** Schematic overview of labeling of HPV pseudovirus with AlexaFluor-647 (see 'Materials and Methods' section for more details). **(B)** AlexaFluor-647 labeled HPV (AF647-HPV) run on an SDS-PAGE denaturing gel and imaged on Azure 600 imaging system (Azure Biosciences). Distinct bands for capsid proteins L1 and L2-FLAG are well resolved, fluorescent, and run at expected molecular weights of 55 and 70 kDa, respectively. **(C)** Representative negative stain electron microscopy image of fluorescently labeled HPV depicts morphologically intact particles. Scale bar: 100 nm. **(D)** Representative immunoblot (top) and quantification (bottom) of reporter GFP expression of HPV-infected HeLa cells, prepared using standard procedures (*in vitro*, MOI=1) or fluorescent labeling procedures (AF647-HPV, MOI=1). At 48 h.p.i., cells were harvested, lysed, and the resulting whole-cell lysate was subjected to SDS-PAGE, followed by immunoblotting with antibodies recognizing GFP and β-actin as a loading control. GFP band intensities are normalized to β-actin in each sample. Total GFP band intensities per condition are normalized against the *in vitro* sample to give normalized infectivity (Normalized inf., %) (Welch's t-test) (N = 3). **(E)** Immunoblots and quantification of GFP expression during AF647-HPV infection (MOI=5) in the presence of γ-secretase inhibitor (XXI), Arf1 nucleotide exchange inhibitor (NAV-2729), or CDK1 inhibitor (RO-3306) were performed as in (D). Cells were either treated with Dimethyl sulfoxide (DMSO, mock control), 2 μM XXI, 25 μM NAV-2729, or 9 μM RO-3306. For quantification, DMSO was used as the normalization standard. (Welch's t-test, **** p<0.0001, *** p= 0.0009, *** p= 0.0001) (N=3). **(F)** Representative confocal fluorescence microscopy images of HA-COPγ1-mCherry transfected and AF647-HPV-infected HeLa cells. HeLa cells were transfected for 24 hours and then infected at MOI=5 for 22 hours post-infection (h.p.i). Images are pseudo-colored as follows: Nucleus (DAPI, blue), HPV16L1 (capsid protein L1 antibody, green), COPγ1 (HA-COPγ1-mCherry plasmid, red), and AF647-HPV (AF647-HPV, magenta). Scale bar: 20 μm **(G, H, I)** Quantification of Pearson Correlation Coefficient (PCC) for fluorescence overlap between (G) capsid protein L1 and HA-COPγ1-mCherry (r=0.5482), (H) capsid protein L1 and AF647-HPV (r=0.7846), and (I) AF647-HPV and HA-COPγ1-mCherry (r=0.4711). N=3, n=30 per condition. **(J)** Quantification of Pearson's correlation coefficient (PCC) between the HPV capsid protein L1 and the early endosome marker EEA1 at 4 h.p.i. for unlabeled (r = 0.4478) and fluorescently labeled (r = 0.4043) virus (MOI=5). N=3, n = 30 images per condition. **(K)** Quantification of Pearson's correlation coefficient (PCC) between the HPV capsid protein L1 and the trans Golgi network TGN46 at 16 h.p.i. for unlabeled (r = 0.3990) and fluorescently labeled (r = 0.3949) virus (MOI=5). N=3, n = 30 images per condition.

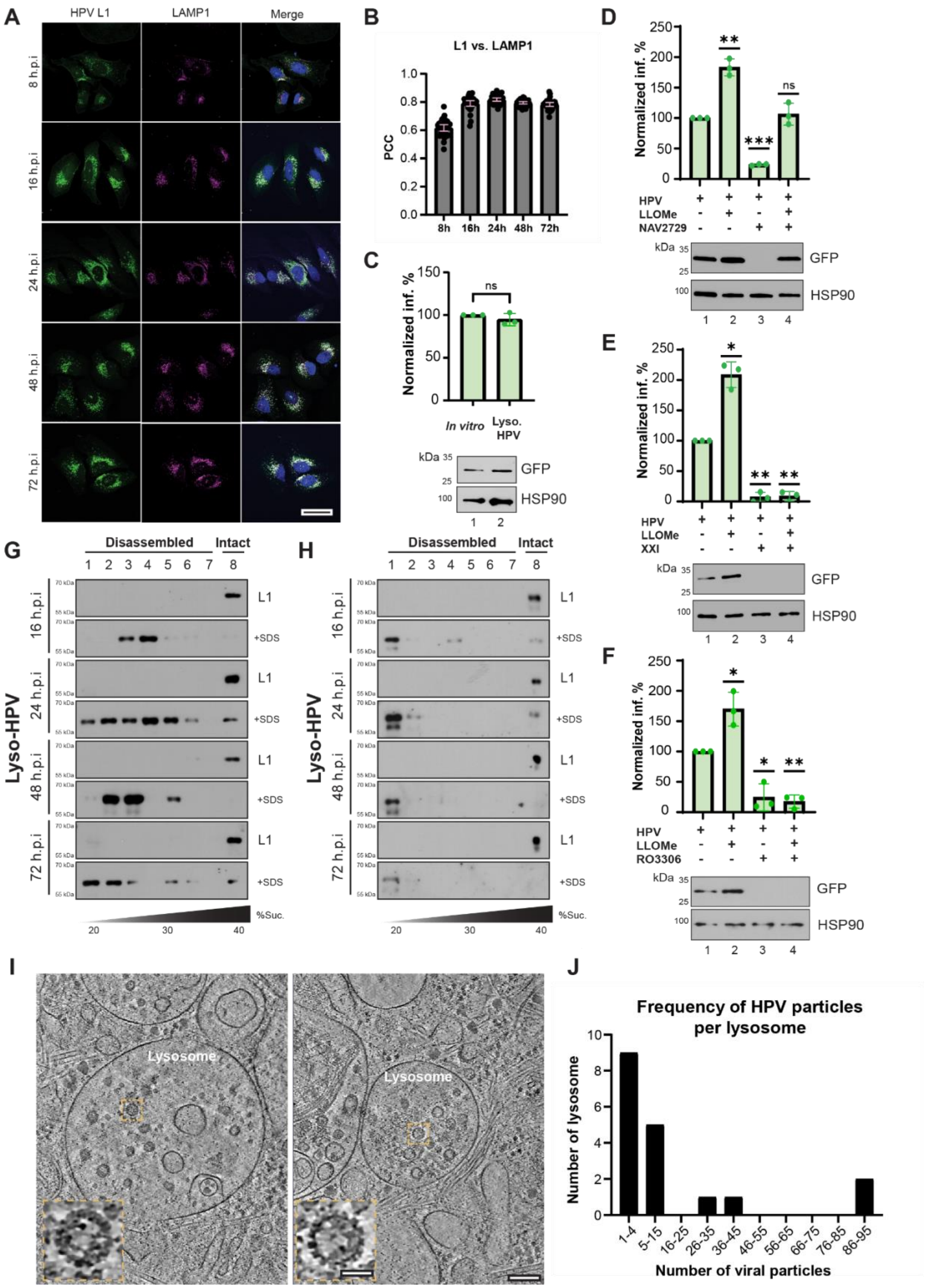
A
HPV L1
LAMP1
Merge
8 h.p.i
16 h.p.i
24 h.p.i
48 h.p.i
72 h.p.i
B
L1 vs. LAMP1
PCC
8h 16h 24h 48h 72h
C
Normalized inf. %
ns
In vitro
Lyso. HPV
kDa
GFP
HSP90
D
Normalized inf. %
**
***
ns
HPV
LLOMe
NAV2729
GFP
HSP90
E
Normalized inf. %
*
**
HPV
LLOMe
XXI
GFP
HSP90
F
Normalized inf. %
*
**
HPV
LLOMe
RO3306
GFP
HSP90
G
Disassembled
Intact
Lyso-HPV
16 h.p.i
24 h.p.i
48 h.p.i
72 h.p.i
L1
+SDS
%Suc.
H
Disassembled
Intact
Lyso-HPV
L1
+SDS
%Suc.
I
Lysosome
J
Frequency of HPV particles per lysosome
Number of lysosome
Number of viral particles
1-4
5-15
16-25
26-35
36-45
46-55
56-65
66-75
76-85
86-95

**Figure S2. HPV persists in host lysosomes for an extended period of time. (A)** Representative confocal fluorescence microscopy images of infected HeLa cells (MOI=10) at 8-, 16-, 24-, 48-, and 72-hours post-infection (h.p.i). Images are pseudo-colored as follows: Nucleus (DAPI, blue), HPV (capsid protein L1, green), and lysosomes (LAMP1, magenta). Scale bar: 20 µm **(B)** Quantification of Pearson Correlation Coefficient (PCC) for fluorescence overlap between capsid protein L1 and lysosomal marker LAMP1 at 8, 16, 24, 48, and 72 h.p.i (r=0.6169, r=0.79, r=0.817, r=0.7946, r=0.7833). N=3, n=30 per condition. **(C)** Representative immunoblot (bottom) and quantification (top) of reporter GFP expression in HPV-infected HeLa cells (MOI=~1), prepared using standard procedures (*in vitro*) or extracted from lysosomes (Lyso. HPV). The amount of added virus was standardized based on GFP plasmid levels determined by PCR (see Materials and Methods for details). At 48 h.p.i., cells were harvested, lysed, and the resulting whole-cell lysate was subjected to SDS-PAGE, followed by immunoblotting with antibodies against GFP and HSP90, with the latter used as a loading control. GFP band intensities are normalized to HSP90 in each sample. Total GFP band intensities per condition are normalized against the *in vitro* sample to give normalized infectivity (Normalized inf., %) (Welch's t-test) (N = 3). **(D)** Top: Representative immunoblots of HPV-infected HeLa cells (MOI=10) treated with DMSO, LLOMe, NAV-2729, or both LLOMe and NAV-2729. NAV-2729 (20 µM) was added at the time of infection, whereas LLOMe (1 mM) was added at 16 h.p.i. for 30 min. At 48 h.p.i., cells were lysed and analyzed by SDS–PAGE and immunoblotting with antibodies against GFP and HSP90, which served as a loading control. Bottom: Quantification of GFP signal intensities normalized to HSP90 and expressed relative to the DMSO-treated control. (Welch's t-test,** p=0.009, *** p= 0.0001, ns p= 0.6065) (N=3) **(E)** Top: Representative immunoblots of HPV-infected HeLa cells (MOI=10) treated with DMSO, LLOMe, XXI, or both LLOMe and XXI. XXI (1 µM) was added at the time of infection, whereas LLOMe (1 mM) was added at 16 h.p.i. for 30 min. At 48 h.p.i., cells were lysed and analyzed by SDS–PAGE and immunoblotting with antibodies against GFP and HSP90, which served as a loading control. Bottom: Quantification of GFP signal intensities normalized to HSP90 and expressed relative to the DMSO-treated control. (Welch's t-test,* p=0.0124, ** p= 0.0024, ** p= 0.0025) (N=3) **(F)** Top: Representative immunoblots of HPV-infected HeLa cells (MOI=10) treated with DMSO, LLOMe, RO3306, or both LLOMe and RO3306. RO3306 (9 µM) was added at the time of infection, whereas LLOMe (1 mM) was added at 16 h.p.i. for 30 min. At 48 h.p.i., cells were lysed and analyzed by SDS–PAGE and immunoblotting with antibodies against GFP and HSP90, which served as a loading control. Bottom: Quantification of GFP signal intensities normalized to HSP90 and expressed relative to the DMSO-treated control. (Welch's t-test,* p=0.0494, * p= 0.0279, ** p= 0.0058) (N=3) **(G, H)** Lysosome-extracted HPV (MOI=50) collected at the indicated time points from TMEM192-FLAG–expressing HEK293 (G) or HaCaT (H) cells pretreated with 9 µM RO-3306 was layered onto a discontinuous 20%–40% sucrose gradient and subjected to centrifugation. Fractions were collected from the top of the gradient, and L1 distribution was assessed by immunoblotting. L1 detected in fractions 1–7 represents disassembled viral material, whereas L1 in fraction 8 corresponds to intact virions. **(I)** Slices through two different tomograms showing lysosome and distinct HPV in the inset. Scale bar: 100 nm, 25 nm (inset). **(J)** Frequency distribution of HPV particles within lysosomes. Among the 18 lysosomes analyzed, 9 contained 1–4 HPV particles, 5 contained 5–15 particles, 1 contained 33 particles, 1 contained 43 particles, and 2 contained 86–

95 particles. We identified these lysosomes across 14 tomograms and 11 lamellae from 11 individual cells and four biological replicates.

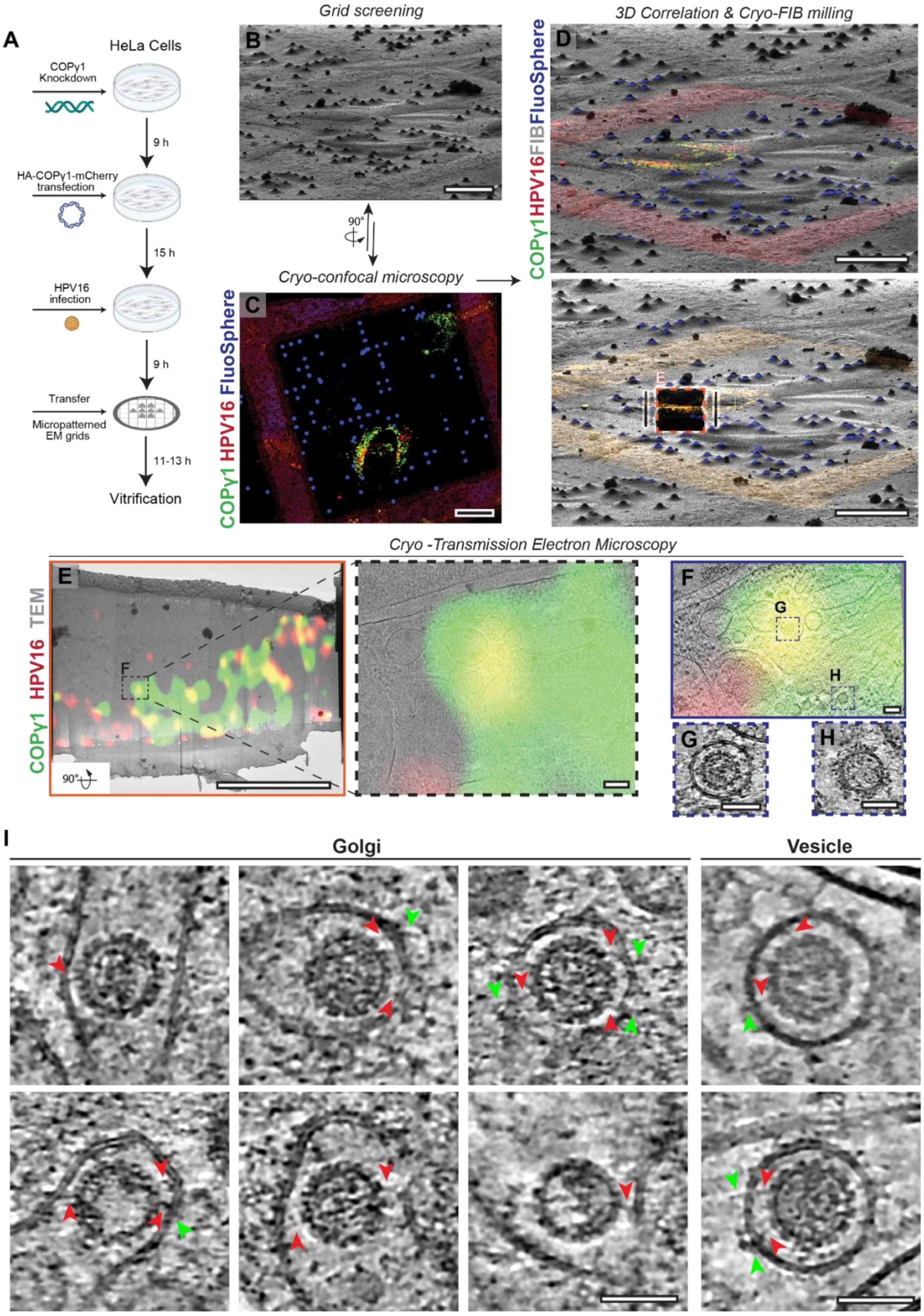
A
HeLa Cells
COPγ1 Knockdown
9 h
HA-COPγ1-mCherry transfection
15 h
HPV16 infection
9 h
Transfer
Micropatterned EM grids
11-13 h
Vitrification
Grid screening
B
90°
Cryo-confocal microscopy
C
COPγ1 HPV16 FluoSphere
3D Correlation & Cryo-FIB milling
D
COPγ1HPV16FIBFluoSphere
E
Cryo -Transmission Electron Microscopy
E
COPγ1 HPV16 TEM
F
90°
F
G
H
G
H
I
Golgi
Vesicle

**Figure S3.** ***In situ* cryo-electron tomography workflow for visualization of HPV trafficking.** **(A)** Schematic of HPV infected HeLa cells (MOI=100) on micropatterned electron microscopy grids (see 'Materials and Methods' section for details). The grids are manually checked under a widefield microscope for quality and subsequently plunge-frozen in liquid ethane. FluoSpheres (diameter – 2 µm) are added immediately before plunge freezing. **(B)** Representative cryo-focused ion beam (cryo-FIB) image of a vitrified grid. Scale bar: 20 µm. **(C)** Maximum Intensity projection of a z-stack obtained from cryo-confocal fluorescence microscopy of the grid square shown in (B). Image is pseudo-colored as follows: HA-COPγ1 (green), 2 µm FluoSpheres (blue), and AlexaFluor-647 labeled HPV (red). Scale bar: 20 µm. **(D)** Overlay of the ion beam image in (B) with the cryo-fluorescence microscopy image in (C) for precise 3D correlation. Top: before cryo-FIB milling; bottom: post cryo-FIB milling. Scale bar: 20 µm. **(E)** Left: Overlay of the cryo-fluorescence image with signals for HA-COPγ1 (green) and HPV (red) on a medium magnification cryo-transmission electron microscopy image (6500×). The black dashed rectangle highlights the region selected for tilt series acquisition. Right: Zoomed-in image of the fluorescence overlay on the cryo-TEM image. Scale bars: 5 µm (lamella), 100 nm (zoom). **(F)** Slice through a reconstructed tomogram acquired at the region in (E) overlaid with cryo-confocal fluorescence data. Note: The slice is presented is rotated by 90° in comparison to the lamella view (E). Scale bar: 100 nm. **(G)** Representative slice through a tomogram, at high fluorescence overlap, highlighting a COPI-coated vesicle containing HPV. Scale bar: 50 nm. **(H)** Representative slice through a tomogram, with no fluorescence for HPV, showing a COPI-coated vesicle without an HPV particle. Scale bar: 50 nm. **(I)** Gallery of representative slices through tomograms showing HPV particles within Golgi compartments (left) or vesicles (right). Red arrowheads indicate proteinaceous densities bridging the viral capsid and the membrane. Bright green arrowheads indicate extension of these bridging densities outside the Golgi lumen Scale bars: 50 nm.

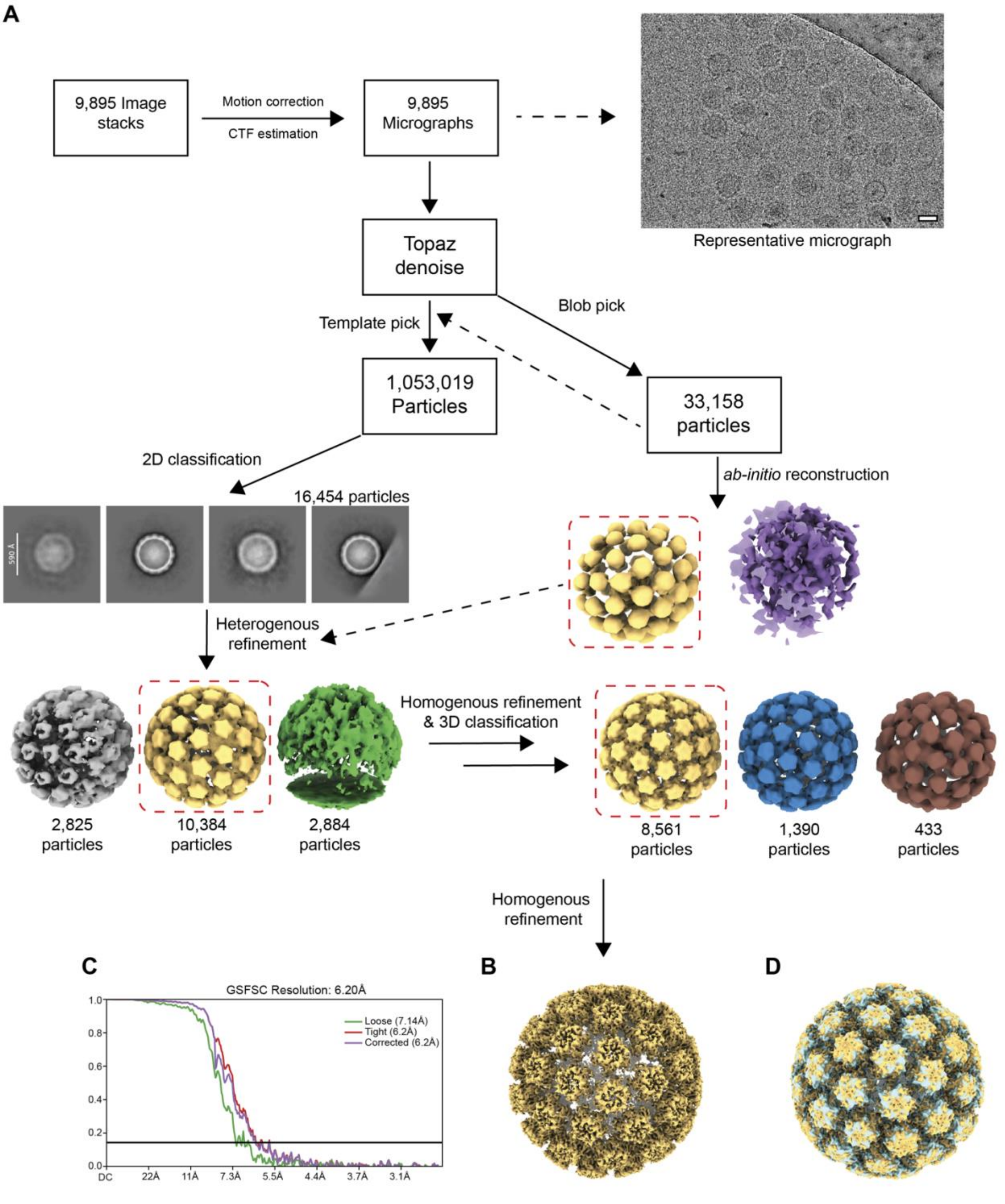


**Figure S4. Cryo-EM single particle analysis workflow for lysosomal HPV dataset. (A)** Overview of processing pipeline. All processing steps were performed using cryoSPARC v4.7.1. A representative micrograph containing HPV purified from lysosomes is depicted. Micrographs were denoised using TOPAZ to aid with particle picking. 700 particles were manually picked from a subset of 144 micrographs to generate 2D templates for template-based picking (dashed lines)

and to generate a low-resolution *ab initio* reconstruction to be used as input volume for heterogeneous refinement (dashed line). 1,053,019 and 33,158 particles were picked using a template picker and a blob picker, respectively. Particles were independently cleaned and sorted through subsequent 2D classification, heterogeneous refinement, and 3D classification, resulting in a final particle stack of 8,561 particles. These particles were re-extracted and refined, initially with C1 symmetry and finally with icosahedral symmetry. **(B)** Final map of lysosomal HPV at 6.20 Å. **(C)** Gold Standard Fourier Shell Correlation (GSFSC). **(D)** Overlay of the cryo-EM maps of the HPV purified from lysosome (yellow) and previously determined structure of HPV at pH 7.4 (EMD – 23081, cyan). HPV purified from lysosomes aligns perfectly well with the previously determined structure of HPV, including the hexavalent and pentavalent subunits.

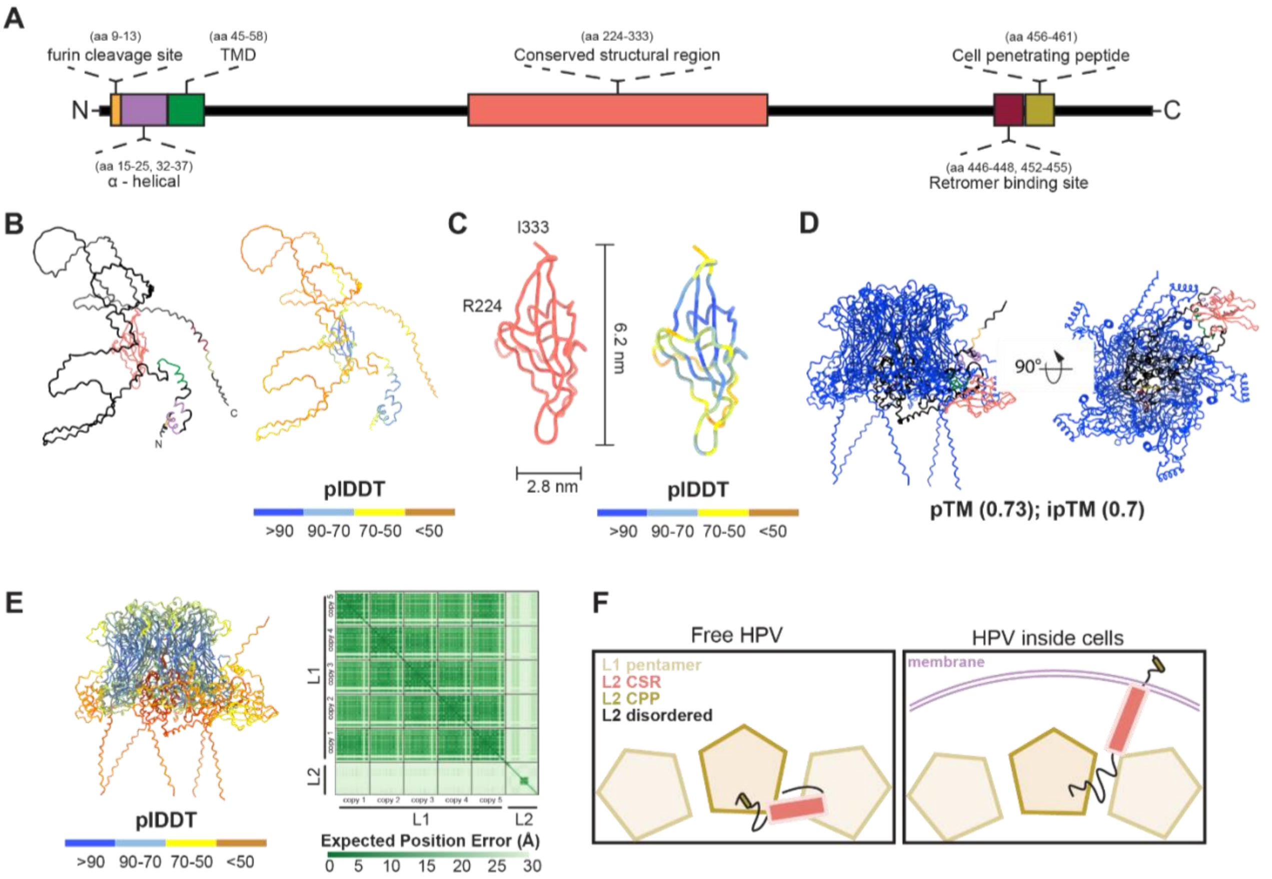


**Figure S5. Structural modeling of L2 and L1-L2 complex using AlphaFold3.** (**A**) Cartoon schematic of HPV capsid protein L2 with key regions highlighted as follows: Furin cleavage site (orange), α-helical (lilac), transmembrane domain (green), central structured region (salmon pink), retromer binding site (RBS, maroon), cell penetrating peptide (CPP, dark yellow), and unstructured/disordered region (black). (**B**) Left: Structural prediction of HPV L2 with domains colored, as described in (A). All predictions were performed on the AlphaFold3 server [133]. Right: Structural model of HPV L2 colored per pLDDT score. The pLDDT > 90 (dark blue) indicates high estimated accuracy of backbone and side chain rotamers, whereas pLDDT > 70 (light blue) indicates confident backbone prediction. The central structured region (CSR, amino acids 224-333) is predicted with high confidence. (**C**) Left: AlphaFold3 prediction of the CSR alone with dimensions denoted. Right: Structural model of HPV L2 CSR colored per pLDDT score. Within the CSR, some regions are predicted with low confidence. (**D**) Structural prediction of full-length HPV capsid L1 protein (5 copies) with full-length L2 protein complex. All L1 chains are colored in blue. L2 is colored as in (B). The predicted template modeling score (pTM) and the interface predicted template modeling score (ipTM) are 0.73 and 0.7, respectively. An ipTM score between 0.6 and 0.8 is a gray zone of prediction. In contrast, a pTM score of 0.73 is good (values greater than 0.5 are considered reliable), indicating overall high confidence in the predicted fold of the complex. (**E**) Left: Structural prediction of full-length L1 pentamer – full-length L2 co-complex, colored based on the pLDDT score. Right: Heatmap showing expected position error (in Ångströms) colored by pLDDT score. **(F)** Cartoon schematic of L2 organization relative to the L1 capsid in free HPV and HPV within membranes. Note: the sizes are not to scale.

### Materials and methods

#### Cell Culture

HeLa cells, used for most experiments, were obtained from the American Type Culture Collection (ATCC, CCL-2). Human Embryonic Kidney (HEK) 293 cells were obtained from ATCC (CRL-15730. HaCaT cells were purchased from AddexBio Technologies. HEK293TT cells, used for HPV pseudovirus production, were provided by Dr. Christopher Buck from the National Cancer Institute in Rockville, MD. All cell lines were cultured in Dulbecco's Modified Eagle's Medium (DMEM) (Gibco, #12430-054) supplemented with 10% fetal bovine serum (FBS) (R&D System, #S11150) and 1× penicillin-streptomycin (Invitrogen, #15140148), referred to as complete media. Cultures were maintained at 37°C with 5% $CO_2$.

#### Production of HPV pseudovirus

We used previously published protocols for the production of HPV pseudovirus [73,76,77]. Briefly, p16sheLL (a gift from Dr. John Schiller, National Cancer Institute, Rockville, MD; also available on Addgene, #37320) was modified for the incorporation of a 3× FLAG tag to the C-terminus of the L2, designated as p16sheLL.L2F [81]. HEK293TT cells were co-transfected with p16sheLL.L2F and a GFP reporter plasmid (pcDNA3.1) [43] using polyethyleneimine (PEI; Polysciences Inc.). After 48 hours, cells were lysed in a buffer containing 0.5% Triton X-100, 10 mM $MgCl_2$, and 5 mM $CaCl_2$. The lysate was then incubated overnight at 37°C with 250 U/mL Benzonase Nuclease (Sigma, #E1014-25KU) and 10 U/mL Plasmid-Safe DNase (Lucigen, #NC1424105). Following incubation, the extract was subjected to ultracentrifugation on an OptiPrep (Milipore Sigma, #D1556) gradient (27%, 33%, and 39%) at 45,000 rpm for 4 hours at 16°C using a Beckman SW55 Ti rotor. The purity of the pseudovirus preparation was assessed by SDS-PAGE followed by Coomassie staining (Invitrogen, LC6065). The pseudovirus titer was estimated by quantifying GFP expression in HeLa cells 48 hours post-infection using flow cytometry.

#### Generation of fluorescently labeled pseudovirus

The HPV pseudovirus from the step above was concentrated to ~1 mg/mL using an Amicon Ultra Centrifugal Filter with a 100 kDa molecular weight cut-off (MWCO) (Millipore Sigma, #UFC9100) by centrifuging at 4000 rpm for 10 minutes at 16°C. For the labeling reaction, the pH of the concentrated virus was adjusted by diluting it (1:10) in 1 M sodium bicarbonate buffer (pH 8.3). Subsequently, 1 µL of 10 mg/mL Alexa Fluor 647 NHS Ester (Thermo Fisher, #A20006) was added, while vortexing, to achieve a 1:1 molar ratio of protein to dye. The mixture was incubated at room temperature for 1 hour with continuous agitation. Separately, a Sephadex G-25 filter column (Cytiva, #28918004) was equilibrated by centrifuging twice with PBS at 4000 rpm for 10 seconds each time. The virus-dye mixture suspension was adjusted to a final volume of 500 µL with DPBS/0.8 M salt buffer (1× DPBS, 762 mM NaCl, 0.9 mM $CaCl_2$, 0.5 mM $MgCl_2$, 4.8 mM KCl). The entire mixture was loaded onto the pre-equilibrated G-25 column and centrifuged for 2 seconds per cycle to collect the fractions. This step was repeated until all the volume had been processed. Blue resin indicates the presence of free dye. Note: It is essential to avoid repeated freeze-thaw cycles of the HPV pseudovirus, as it compromises the virus or impairs labeling efficiency. To confirm labeling efficiency, the fluorescently labeled virus was denatured at 90°C for 10 minutes, separated by SDS-PAGE, and imaged on an Azure 600 imaging system (Azure

Biosciences) using AlexaFluor647 settings. Infectivity was assessed as described in the “Immunoblotting” section below. Structural integrity of the capsids was verified according to the “Negative Stain Electron Microscopy” section, and the localization of the fluorescently labeled virus was determined as outlined in the “Immunofluorescence Microscopy” section.

**Transfection and small-interfering RNA (siRNA) knockdown**

For the room temperature confocal microscopy and the cryo-correlative experiments, HeLa cells were seeded at a density of 2.1 × $10^5$ cells per well in a six-well plate (Fisher, #FB012927) and transfected with 1 nM COPγ1 siRNA (5’-CUUGUAAUCUGGAUCUGGA-3’) using Lipofectamine RNAiMAX (Invitrogen, #13778150) in OPTI-MEM (Gibco, #31985062) for 9 hours, with a 1:4 dilution of DMEM, per manufacturer’s instructions. Following this, the cells were washed three times with PBS and transfected with 1 µg of the pCMVTNT-HA-COPγ1-mCherry plasmid for 16-18 hours using the FuGENE HD transfection reagent (Promega, #E2311) in OPTI-MEM with a 1:4 dilution of DMEM, maintaining a 1:3 ratio of plasmid to transfection reagent according to the manufacturer’s guidelines.

For the immunoprecipitation of lysosomes (Lyso-IP) experiment, HeLa cells were seeded in a 15 cm plate (Fisher, #FB012924) and transfected with 5 µg of the TMEM192-HA plasmid using polyethyleneimine (PEI) (Polysciences Inc., #24765) for 24 hours. Following this transfection period, the cells were infected with HPV pseudovirus for 16 hours. Subsequent steps are outlined in the ‘Lyso-IP’ section below.

**Drug treatments**

1.1 × $10^5$ cells per well in a twelve-well plate (Fisher, #FB012928) were treated with either 1 or 2 µM XXI (Millipore Sigma, Cat #565790), 20 or 25 µM NAV-2729 (Millipore Sigma, Cat #SML2238), or 9 µM RO3306 (Selleckchem, Cat #S7747) simultaneously with the addition of the virus. Forty-eight hours post-infection and treatment, the cells were harvested for immunoblot analysis.

For the **L**-Leucyl-**L**-Leucine methyl ester hydrobromide (LLOME) experiment, 16 h.p.i HeLa cells were treated with 1 mM LLOME (Milipore Sigma, #L7393) and incubated for 30 minutes. Following incubation, the cells were washed three times with PBS, then replenished with pre-warmed complete media, and incubated for an additional 32 hours before being collected for immunoblot analysis, as detailed below. Concentrated stock solutions of LLOME were prepared in DMSO (Milipore Sigma, #D2438-50ML) and stored at -80°C.

For the control conditions, an equivalent volume of DMSO was added.

**Immunofluorescence microscopy**

1.1 × $10^5$ HeLa cells were seeded onto glass coverslips (Fisher, #50-189-7777) in a 12-well plate. Post attachment, cells were transfected (where indicated) and/or infected with either unlabeled or fluorescently labeled virus. Cells were fixed at the specified time points using 4% paraformaldehyde (PFA) for 15 minutes, then permeabilized and blocked with 0.2% Triton X-100 in complete media for 1 hour at room temperature. Subsequently, the cells were incubated overnight with the primary antibody, followed by three washes with PBS for 10 minutes each. Next, the samples were incubated with Alexa Fluor–conjugated secondary antibodies, diluted

1:2000 in 0.2% Triton X-100/complete media, for 1 hour at room temperature. Coverslips were mounted with a DAPI-containing mounting medium (Abcam, ab104139) and imaged using a Zeiss LSM 800 confocal laser scanning microscope with a Plan-Apochromat 40×/1.4 oil differential interference contrast M27 objective.

**Immunoblotting**

Forty-eight hours post-infection, cells were washed three times with PBS and lysed in HN buffer [1% Triton X-100, 50 mM HEPES, 150 mM NaCl, 1 mM phenylmethylsulfonyl fluoride (PMSF)], for 10 minutes on ice. The lysate was centrifuged at 16,100*g* for 10 minutes. Following this, the supernatant was collected and mixed with SDS sample buffer containing 5% 2-mercaptoethanol (Milipore Sigma, #M6250). The mixture was then denatured at 90°C for 10 minutes, separated by SDS-PAGE, and transferred to nitrocellulose membranes (Milipore Sigma, #10600003). Membranes were incubated overnight at 4°C with primary antibodies (see Supplementary Table S1) diluted in 1x Tris-buffered saline (TBS) containing 3% skim milk and 0.2% Tween-20. Blots were washed three times for 10 minutes each with TBS containing 0.2% Tween-20 (TBT). The membranes were then incubated with horseradish peroxidase (HRP)-conjugated secondary antibodies in TBT with 3% skim milk at room temperature for 1-2 hours, followed by three 10-minute washes with TBT. Chemiluminescence signals were generated using HRP substrates and detected by exposure to X-ray films (Fisher, # NC9556985).

**Patient sample collection**

Healthy women and patients with known HPV16-positive screening results and diagnoses of cervical intraepithelial neoplasia grade 1 or 2 (CIN1/2) were recruited for this study. Their samples were taken from routine cervical cancer screening using primary HPV testing with the Roche Cobas 4800 HPV detection platform. All samples were taken directly from the cervix, placed in ThinPrep preservative, and sent to the lab for HPV genotyping. Institutional approval to reuse patient samples after clinical care enabled us to use them in our laboratory for our study. Institutional Review Board (IRB)- approved protocols at the University of Michigan are available under ID HUM00282032. All samples are coded, and personal health information is removed (de-identified) before they are shared with potential investigators.

**Immunofluorescence staining of ThinPrep patient samples**

Coverslips were coated to ensure optimal cell adhesion. A 0.1 mg/mL poly-L-lysine (PLL) solution (Sigma-Aldrich, #P6282) was prepared and used to fully cover each coverslip. The coverslips were incubated in the coating solution for 60 minutes at room temperature. After incubation, the coverslips were washed three times with sterile distilled water to remove any unbound polylysine, then allowed to air-dry thoroughly in a sterile environment.

For fixation and attachment of ThinPrep cells to coverslips, the cell sample was gently resuspended by flicking the vial, then transferred to a 15 mL conical tube. The cells were pelleted by centrifugation at 1000 × *g* for 5 minutes. The supernatant was carefully removed, and the pellet was washed with 1× PBS to eliminate the PreservCyt solution. After a second centrifugation, PBS was aspirated, and the cells were fixed by adding 4% paraformaldehyde (PFA) in PBS for 15 minutes at room temperature. The cells were gently washed three times in 1× PBS for 10 minutes each, with liquid always added and removed from the side of the well to avoid disturbing the pellet.

The fixed cell pellet was resuspended in a small volume of PBS (100–500 µL) to create a concentrated cell suspension. A volume of 50–100 µL was spotted onto the center of a poly-L-lysine-coated coverslip, placed in a 12-well plate. The cells were allowed to settle and adhere for 30–60 minutes at room temperature. Attachment was confirmed under a phase-contrast microscope to ensure cells remained on the glass.

For permeabilization and immunofluorescence staining, cells were treated with 3% BSA and 0.3% Triton X-100 in PBS for 1 hour. Three washes with PBS were performed, each for 10 minutes. The cells were then incubated overnight at 4°C with primary antibodies diluted to 1:1000 (rabbit anti-LAMP1 and mouse anti-HPV16 L1). After primary antibody incubation, the cells were washed three times with permeabilization medium (0.3% Triton X-100 in PBS) for 10 minutes each. Secondary antibody was applied at a 1:2000 dilution for 1–2 hours at room temperature in the dark. Following secondary incubation, three additional washes with permeabilization medium were performed, each for 10 minutes. Finally, coverslips were mounted for imaging.

**Tagless Lysosome Immunoprecipitation (Lyso-IP)**

ThinPrep patient samples were pooled and centrifuged at 1,200*g* for 5 min. The resulting pellet was washed twice with PBS, centrifuged again, and resuspended in 1 mL KPBS [(36 mM KCl and 10 mM $KH_2PO_4$, pH 7.25, adjusted with KOH, supplemented with 1× aprotinin (MilliporeSigma, #A1153), 1× pepstatin (MilliporeSigma, #P4265), 1× leupeptin (MilliporeSigma, #L2884), and 1 mM PMSF]. Samples were incubated on ice for 30 min and homogenized for 40 strokes using a tight-pestle Dounce homogenizer (DWK Life Sciences, #358003). The homogenate was centrifuged twice at 1,000*g* for 2 min at 4°C. A 50-µL aliquot (5% of the total sample) was retained as the whole-cell lysate input.

The clarified supernatant was incubated with 50 µL of anti-human TMEM192 magnetic beads (ABclonal Science, #A27784) for 30 min with gentle agitation. Bead-bound lysosomes were washed three times with KPBS and resuspended in 50 µL KPBS. The samples were then disrupted on ice using a handheld sonicator at power setting 2 for five cycles of 10 s on and 50 s off. A portion of the resulting material was analyzed by negative-stain electron microscopy, as described below. The remainder was mixed with SDS sample buffer containing 2-mercaptoethanol, denatured at 75°C for 10 min, and analyzed by SDS–PAGE and immunoblotting for HPV and the indicated organelle markers.

**Lysosome immunoprecipitation (Lyso-IP)**

A confluent 15 cm plate of TMEM192-HA-expressing or TMEM192-FLAG-expressing HeLa cells was washed three times with PBS. The cells were then scraped, centrifuged at 1,200*g* for 5 min, and the pellet was resuspended in 1 mL of cold KPBS (supplemented with 1× aprotinin, 1× pepstatin, 1× leupeptin, and 1 mM PMSF). Samples were incubated on ice for 30 min and homogenized for 40 strokes using a tight-pestle Dounce homogenizer. The homogenate was centrifuged twice at 1,000*g* for 2 min at 4°C. A 50-µL aliquot (5% of the total sample) was retained as the whole-cell lysate input. The supernatant, containing cellular organelles, including lysosomes, was incubated with 50 µL of anti-HA magnetic beads (Thermo Fisher, #88836) on a gentle rotator shaker for 5 minutes. Beads with bound lysosomes were washed three times with PBS. Subsequently, the beads were resuspended in 50 µL of KPBS, incubated on ice for 20 minutes, and briefly sonicated using a handheld sonicator at power setting 2 for 5 cycles of 10 s on, and 50 s off to disrupt the lysosomes. The supernatant (referred to as Lyso-IP in the figure) was collected. 2.5 µL of the supernatant was mixed with SDS sample buffer containing 2-mercaptoethanol and denatured at 90°C for 10 minutes. This mixture was separated by SDS-PAGE alongside different dilutions of pseudovirus, then incubated with HPV16 L1 primary antibody in TBT with 3% skim milk for 1 hour at room temperature, followed by HRP-conjugated secondary antibodies in TBT with 3% skim milk for another hour at room temperature.

For genome-based normalization, HPV was isolated from the lysosomal fraction, and the recovered viral material, along with different dilutions of pseudovirus, was analyzed by PCR using GFP-specific primers. The resulting PCR products were quantified to estimate the abundance of packaged GFP-S reporter plasmids, allowing viral inputs to be normalized independently of L1. Following normalization based on either L1 abundance or packaged GFP-S reporter levels, equivalent amounts of lysosome-derived HPV and purified HPV pseudovirus were added to HeLa cells cultured in 12-well plates. 48 hours post-infection, cells and supernatant were harvested and subjected to SDS-PAGE and immunoblotting staining for GFP.

Separately, the remaining supernatant after Lyso-IP was denatured at 65°C for 10 minutes and subjected to SDS-PAGE and immunoblot analysis, as described above, to assess for cross-contamination from other cellular organelles.

**Disassembly assay**

Lysosomal HPV samples (20 µL) obtained by Lyso-IP, as described above, were layered onto discontinuous sucrose gradients consisting of 60 µL each of 20%, 30%, and 40% sucrose, with 20% at the top and 40% at the bottom, in polycarbonate centrifuge tubes (Beckman Coulter, #343775). Samples were ultracentrifuged at 50,000 rpm for 30 min at 4°C using a Beckman TLA-100 rotor. Eight 25-µL fractions were collected sequentially from the top of each gradient, with fraction 1 representing the top and fraction 8 the bottom. Each fraction was mixed with SDS sample buffer containing 2-mercaptoethanol and denatured at 90°C for 10 minutes, then analyzed by SDS–PAGE followed by immunoblotting.

**Isolation of HPV containing vesicles**

Eight 15 cm plates of HeLa cells infected with unlabeled HPV16L2F pseudovirus (MOI=50 per 15 cm plate) were washed 3 times with ice-cold PBS and then scraped and spun down at 3000 rpm for 10 minutes at 4°C. Subsequently, the pellet was resuspended in 5 mL 10 mM HEPES, pH 7.4, on ice for 15 minutes, following which 5 mL of 2× lysis buffer [0.5M sucrose, 2 mM EDTA, 100 mM HEPES pH 7.4, 2 mM PMSF, 200 mM KCl and 5 mM $Mg(OAc)_2$, Aprotinin 2× (Milipore Sigma, #A1153), Pepstatin 2× (Milipore Sigma, #P4265), Leupeptin 2× (Milipore Sigma, #L2884)] and mechanically homogenized with pre-chilled Dounce homogenizer with a tight-fitting pestle (Kimble "B") on ice (40 strokes). The resulting cell lysate was centrifuged at 1000*g* for 5 minutes at 4°C. Next, the supernatant from the previous step was collected and spun down again at 5000*g* for 10 minutes at 4°C. The supernatant from the second round of centrifugation was collected and incubated with a 300 µL slurry of 'DYKDDDDK' Magnetic Agarose beads (Thermo Fisher, #A36797), which had been pre-washed in 1× lysis buffer, for 1 hour. The beads were then washed 3 times with 1× lysis buffer and the contents eluted with 0.5 mg/mL FLAG peptide (Sino Biological, #PP101274) for 10 minutes at 4 °C with gentle rocking. Immediately following elution, the eluate was used for negative staining and immunogold labeling for anti-M2FLAG (Milipore Sigma, #F3165) as outlined below.

**Negative stain electron microscopy**

Carbon-coated copper grids (300 mesh; Electron Microscopy Sciences, #CF300-CU-50) were glow-discharged for 30 seconds at 10 mA using an EasiGlow system (Pelco). A 3 µL drop of the sample was applied to the grid and allowed to sit for 1 minute at room temperature. Excess liquid was blotted off with Whatman 1 filter paper (Cytiva, #1001090). The grid was washed twice with Milli-Q water and once with 0.75% uranyl formate (Electron Microscopy Sciences, # 16984-59-1). The grid was stained for 1 minute in a fresh droplet of 0.75% uranyl formate before being subjected to final manual blotting and air drying. The grids were stored until further use.

Grids were imaged using a Morgagni transmission electron microscope (Thermo Fisher Scientific) at an acceleration voltage of 100 kV. Images were acquired using a Gatan Orius SC200 CCD camera at a pixel size of 2.1 Å (corresponding to a nominal magnification of 22,000×) using Digital Micrograph software (Gatan Inc).

**Immunogold labeling of HPV-containing vesicles**

Carbon-coated copper grids (300 mesh) were glow-discharged as described above. 3µL of the FLAG-IP eluate was applied to the grids. After 1 min, excessive liquid was removed by manual blotting using Whatman 1 filter paper. Next, a blocking buffer (PBS, pH 7.4, 0.1% w/v cold water fish skin gelatin (Aurion, #CFWS Gelatin)) was applied to the grids for 10 minutes, followed by blotting of the excess buffer with filter paper. Subsequently, M2-FLAG antibody diluted 1:10 in blocking buffer was applied to the grids and incubated for 1 hour at room temperature. Following this, grids were manually blotted and washed three times with blocking buffer and twice with 1× PBS, with excess solution blotted off between each wash. Next, 12 nm Colloidal Gold AffiniPure Donkey Anti-Mouse IgG (Jackson ImmunoResearch Laboratories, #715-205-150) was diluted 1:4 in blocking buffer and applied to the grids and incubated for 1 hour. Following this, the grids were washed five times with 1× PBS and three times with Milli-Q water, and excess liquid was manually blotted away between washes. Finally, grids were stained with 3 µL of 0.75% uranyl formate for 5 minutes, as described above. The grids were allowed to dry and imaged on the transmission electron microscope (T12, Tecnai) operated at an acceleration voltage of 120kV. The images were recorded on a Gatan Rio9 CMOS Detector with a pixel size of 2.41 Å, using SerialEM [134].

**Sample preparation and data collection for cryo-electron microscopy (cryo-EM)**

For cryo-EM sample preparation, a similar lyso-IP protocol above was followed, with the exception that 8, 15 $cm^2$ plates were used and the pellet was resuspended in 8 mL of cold PBS. Beads with bound lysosomes were washed three times with PBS. Subsequently, the beads were resuspended in 50 µL of 0.2 M Sodium Acetate/Acetic acid at pH 4.5, incubated on ice for 20 minutes, and briefly sonicated using a handheld sonicator at power setting 2 for 5 cycles of 10 s on, and 50 s off to disrupt the lysosomes. Lyso-IP fraction was centrifuged down at 5,000*g* for 10 minutes at 4°C, and subsequently concentrated using an Amicon ultra centrifugal filter with a 100 kDa molecular weight cut-off (MWCO) by centrifuging at 4000 rpm for 10 min. at 16°C. The concentrate was checked for quality by negative staining.

R2/2 with 2 nm carbon support 200 mesh grids (Quantifoil) were glow-discharged using PELCO *easiGlow* for 1 minute at 15 mA current. 3 µl of concentrated lysosomal HPV sample was applied to the grids, and blotted for 3 seconds on a Vitrobot (mark IV, ThermoFisher Scientific) operated at 8°C and 100% humidity with blot force 5 and plunge frozen in liquid ethane. Images were acquired using a Titan Krios G4i operated at 300 kV equipped with a bioQuantum energy filter with a slit width of 20 eV and a Gatan K3 detector. Automated collection was performed at nominal magnification of 64,000×, resulting in a calibrated pixel size of 1.37 Å/pix using serialEM v4.2. A total of 50 frames were collected with a cumulative dose of 50.4 $e^{-}$/Å$^2$, and defocus cycling from -3 µm to -1 µm.

**Single particle cryoEM image processing**

All subsequent processing steps were performed using cryoSPARC v4.7.1 [135]. A total of 9,895 movies were aligned, dose-weighted, and summed using full-frame motion correction. Motion corrected micrographs were used to estimate the contrast transfer function using CTFFINFD4 [136]. Topaz was used to denoise the micrograph to aid with particle picking [137]. Initially, 700 particles were manually selected from 144 micrographs to generate a 2D template for template-picking and *ab initio,* which would then be used as an input reference for heterogeneous refinement. Template picking and blob picking yielded 1,053,019 and 33,158 particles from 9,985 micrographs, respectively. The particles were extracted using a box size of 1024 pixels and Fourier-cropped to 256 pixels. The extracted particles underwent multiple rounds of 2D classification to remove most empty classes, followed by joining to eliminate duplicates, resulting in a particle set of 16,454 particles. This particle set was classified to exclude particles at the edge of the hole or on the foil adjacent to it. Heterogeneous refinement and 3D classification were used to further clean these particles, yielding a final set of 8,561 particles that were re-extracted at a box size of 800 pixels. Unbinned particles were initially refined using C1 symmetry and then icosahedral symmetry to obtain the final map of 6.20 Å, as measured by Gold Standard Fourier Shell Correlation (GSFSC). ChimeraX [138,139] was used to visualize and render the maps.

***In situ* cryo-correlative light and electron microscopy (cryo-CLEM)**

<u>Grid micropatterning</u>

Au 200 mesh grids with a holey R1/4 $SiO_2$ film (Quantifoil Micro Tools GmbH, #Q250AR-14S) were glow-discharged for 30 seconds at 5 mA on both sides using an EasiGlow system (Pelco). Following this, grids were placed on a glass slide, secured with PDMS stencils (Alvéole), and coated with 8 µL of poly-L-lysine (Sigma Aldrich, #P6282) for 30. Following the poly-L-lysine coating, the grids were washed four times with Milli-Q water. Next, 8 µL of 100 mg/mL PEG-SVA (Laysan, #MPEG-SVA-5000), diluted in pH 8.5 HEPES buffer (Sigma Aldrich, #P7626), was added to the grids for 1 hour, followed by four additional Milli-Q water washes. The grids were then coated in dark with PLPP gel (Alvéole, #B004) diluted in 100% ethanol at a 1:7 ratio (1 PLPP gel to 7 100% ethanol) for 30 minutes. After coating, the grids were micropatterned using a Leica

DMi8 microscope equipped with an Alvéole PRIMO 2 system, utilizing Leonardo software[140] to create circular patterns with a 50 μm diameter centered on the grid squares at a laser power of 60 mJ/mm². The grids were washed four times with Milli-Q water and stored in the dark at 4 °C until further use. Grids were ensured not to dry out to prevent damage to the film.

Cell culture

2.1 x $10^5$ HeLa cells were seeded per well in a six-well plate and treated as described earlier (see '*Transfection and siRNA knockdown*'). In the meantime, micropatterned grids were sterilized with ethanol and placed in a biosafety cabinet. The grids were transferred to a 35 mm glass-bottom dish (MatTek Life Sciences, #P35G-1.5-14-C) and washed four times with Milli-Q water. Next, the grids were coated with fibronectin for at least 1 hour to promote cell growth on the micropatterned areas [140]. 9 hours post-infection, the cells were washed three times with PBS, trypsinized, and added dropwise onto the fibronectin-coated grids, aiming for approximately 1 to 2 cells per grid square. If the cells clumped or were not evenly distributed, the droplet was replaced with a new one until the desired confluency was achieved. Once satisfactory, the MatTek dish was placed in an incubator at 37°C with 5% $CO_2$ for 5 to 10 minutes to allow cells to settle on the grid film. Finally, 2 mL of fresh, pre-warmed complete media was added to the dish. All procedures were conducted within the biosafety cabinet to maintain sterility. Grids containing HeLa cells were kept in a 37°C incubator until vitrification.

Vitrification

2 μm blue (365/415 nm) FluoroSpheres (Invitrogen, #F8824) were sonicated in a water bath for 10 seconds and then diluted 1:75 in PBS. Immediately before vitrification, 3 μL of the diluted FluoroSpheres were applied to the side of the grids with cells. The grids were back-side blotted with filter paper (Ted Pella, #47000-200), and then plunged-frozen into liquid ethane at -186°C using a GP2 automatic grid plunger (Leica Microsystems). The parameters for blotting were set to a 10-second blot time, with a chamber temperature of 37°C and 90% humidity. The plunge-frozen grids were then clipped into Cryo-FIB Autogrids (Thermo Fisher Scientific, #1205101) with a C-clip (Thermo Fisher Scientific, #1036171) and stored in liquid nitrogen until further use.

For lysosomal cryo-CLEM experiments, all the parameters and subsequent plunging conditions were the same as described above, with the following exception: 50 nM Lysotracker Green (Invitrogen, #L7526) Was added 3-5 minutes before vitrification.

Cryo-confocal fluorescence microscopy

The quality of the grid was assessed in widefield mode on a Stellaris-5 cryo-confocal laser scanning microscope equipped with a cryo-stage and 50×/0.9 numerical aperture objective (Leica Microsystems). The entire grid was mapped in wide-field mode using reflection. Z-stacks of the desired grid square were taken in confocal line scanning mode with either Hoechst34580 (420-450 nm emission), Lysotracker Green (511-558 nm emission), mCherry (597-642 nm emission), and Alexa Fluor 647 (663-750 nm emission) built in settings from LAS X software and 0.3 μm step size.

Cryo-focused ion beam (cryo-FIB) milling

After cryo-fluorescence data acquisition, grids were loaded onto the stage of an Aquilos 2 (Thermo Fisher Scientific) maintained at a temperature of –185°C. To reduce charge-induced movement during cryo-FIB milling and enhance sample conductivity, grids were sputter-coated with an inorganic platinum layer using the following settings: current, 10 mA; pressure, 0.1 mbar; voltage, 1 V; and run time, 20 seconds. Subsequently, a protective layer of organometallic platinum was deposited using a Gas Injection System (GIS) for 30 seconds [141].

Full grid orientation was determined by correlating whole-grid reflection cryo-confocal images with full-grid SEM images using MAPS software (version 3.20), allowing for the identification of the desired grid squares that had been previously imaged. Annotated squares were selected for lamella preparation. For each square, a cryo-FIB image was acquired at 30 pA, 30 kV, 500 ns, 0.7 × 0.5 k resolution, and a magnification of 1,500×. The 3D Correlation Toolbox (3DCT) was then used to align confocal z-stacks and ion beam images, using the position of the FluoSpheres to precisely position lamella milling sites [142].

First tension-relief trenches were milled using a beam current of 1 nA [143]. Next, the cellular regions of interest were sequentially thinned down to 5 µm, 3 µm, and 1 µm using currents of 1nA, 500 pA, and 300 pA, respectively, to generate ‘rough’ lamellae. Throughout the milling process, SEM imaging was used to monitor milling progress and lamella integrity. This workflow was repeated for each target square, capturing ion beam images and correlating as described.

After all desired rough lamellae were generated, each lamella was polished to a final thickness of ~200 nm using 50 pA beam current. Following fine milling, cryo-FIB images with the same settings were acquired for all squares, and SEM images centered on the lamellae were captured at magnifications of 1,200× and 5,000×. The final step involved a second round of correlation using 3DCT to align post-milling fluorescence signals on the lamellae [144].

Tilt series data acquisition and tomogram reconstruction

Cryo-FIB milled grids with lamellae were imaged on a 300 kV Titan Krios G4i transmission electron microscope (Thermo Fisher Scientific) equipped with a K3 direct electron detector and a Gatan imaging filter, operated in counting mode. Dose-symmetric tilt series were acquired at correlated regions using SerialEM s, totaling 37 tilts per area of acquisition, beginning at the pre-tilt angle and an increment of 3°. Tilts were recorded at a magnification corresponding to a pixel size of 2.654 Å/pixel, with a total electron dose of 160 $e^-/Å^2$ per tilt series and a 20 eV slit width. The tilt series were aligned using patch tracking and reconstructed by weighted back-projection in IMOD [145–148]. ‘SIRT-like’ (15 iterations) or Nonlinear Anisotropic Diffusion (NAD) filtering was applied to visually enhance reconstruction quality.

**Segmentation**

4× binned tomograms were used for segmentation with Membrane-Seg [149], employing experimental models v10_beta_FAaug -- model training with Fourier amplitude augmentation. The segmented membranes were then imported into Amira (Thermo Fisher Scientific) for manual refinement. For ribosome and HPV placements, template matching was performed using the ribosome map (EMD-15636) or HPV (EMD 5932) in WARP [150,151], with coordinates cleaned up in CUBE. Microtubules were segmented using the Cylinder Correlation feature in Amira. The final figures were created in UCSF ChimeraX [138,139] by exporting cellular surfaces from Amira as Wavefront.obj files and importing them into ChimeraX. The ArtiaX plugin [152] was used to display and manipulate the tomogram, segmented cellular features, and ribosomes.

**HPV diameter measurements**

All measurements were performed in IMOD. HPV capsid diameter was measured in the central tomographic slice, defined as the slice containing the largest particle cross-section, and calculated as the maximum distance between opposing L1 pentamers. Vesicle diameter was measured similarly in the slice containing the largest vesicle cross-section and defined as the length of its longest axis. Bridging-density length was defined as the distance from the viral surface to the inner leaflet of the adjacent membrane, whereas density width was defined as its maximum lateral span.

**Statistical analysis**

Immunoblot images were analyzed using FIJI software [153] to quantify GFP levels, as previously described. Briefly, ROI were drawn around each band, and the integrated intensity was measured [43,48,58,60,61,90]. GFP band intensities were normalized to the corresponding actin bands. All statistical analyses were performed using GraphPad Prism 10.4.1.
